# Transcriptome responses to reduced dopamine in the Substantia Nigra Pars Compacta reveals a potential protective role for dopamine

**DOI:** 10.1101/468330

**Authors:** M. Koltun, K. Cichewicz, J.T. Gibbs, M. Darvas, J. Hirsh

## Abstract

Parkinson’s Disease (PD), is a neurodegenerative disorder affecting both cognitive and motor functions. It is characterized by decreased brain dopamine (DA) and a selective and progressive loss of dopaminergic neurons in the substantia nigra pars compacta (SNc), whereas dopaminergic neurons in the ventral tegmental area (VTA) show reduced vulnerability. The majority of animal models of PD are genetic lesion or neurotoxin exposure models that lead to death of dopaminergic neurons. Here we use a DAT:TH KO mouse model that by inactivation of the tyrosine hydroxylase (*Th*) gene in dopamine transporter-expressing neurons, causes selective depletion of striatal dopamine without affecting DA neuron survival. We analyzed transcriptome responses to decreased DA in both pre- and post-synaptic dopaminergic brain regions of DAT:TH KO animals. We detected only few differentially expressed genes in the post-synaptic regions as a function of DA deficiency. This suggests that the broad striatal transcriptional changes in neurodegeneration-based PD models are not direct effects of DA depletion, but are rather a result of DA neuronal death. However, we find a number of dopaminergic genes differentially expressed in SNc, and to a lesser extent in VTA, as a function of DA deficiency, providing evidence for a DA-dependent feedback loop. Of particular interest, expression of *Nr4a2*, a crucial transcription factor maintaining DA neuron identity, is significantly decreased in SNc, but not VTA, of DAT:TH KO mice, implying a potential protective role for DA in the SNc.

## Introduction

Midbrain dopamine neurons in the ventral tegmental area (VTA) and substantia nigra pars compacta (SNc) are the major source of DA in the brain through their projections to the striatum, prefrontal cortex (PFC) and other limbic areas including the amygdala and hippocampus (Björklund and Dunnett 2007). Via these projections to the striatum and PFC, DA neurons play crucial roles in reward, motor and cognitive processes (Wise 2004). SNc and VTA DA neurons have different major projection fields: SNc DA neurons project primarily to the dorsal striatum (DS) whereas VTA DA neurons project to the ventral striatum (VS) and to the PFC, hippocampus and amygdala. Loss-of-function studies using neurotoxins and genetic approaches indicate that VTA and SNc DA neurons have specific involvement in reward, motor, memory and learning processes (Darvas, Wunsch, et al. 2014; Darvas, Henschen, and Palmiter 2014; Darvas and Palmiter 2009, 2011; Baunez and Robbins 1999; Chudasama and Robbins 2006; Da Cunha et al. 2002; De Leonibus et al. 2007; Mura and Feldon 2003). Parkinson’s disease (PD) is characterized by early degeneration of SNc neurons and the presence of alpha-synuclein inclusion bodies (Lewy bodies), yet only much later is there involvement of the VTA in PD. Clinical observations in PD patients suggest that loss of SNc DA neurons contribute to both motor and cognitive symptoms of PD (R Cools et al. 2001; Roshan Cools et al. 2003; Sawamoto et al. 2008).

The earliest changes induced by alpha synuclein accumulation are predegenerative loss of DA neurotransmission, followed by axonopathy and subsequent loss of DA neurons (Luk et al. 2012; Lundblad et al. 2012). Early changes in midbrain neurons resulting from loss of DA may be involved in the chain of pathologic processes leading to degeneration of DA neurons. We utilize a genetic model that allows investigation of the effects of DA loss on midbrain gene expression without degeneration of midbrain DA neurons. This model lacks tyrosine hydroxylase (TH) selectively in DA-transporter (DAT) expressing neurons, resulting from a conditional (floxed) *Th* deletion caused by expression of Cre recombinase in DAT neurons (Henschen, Palmiter, and Darvas 2013; Jackson et al. 2012; Zhuang et al. 2005). Although these animals have severe loss of striatal DA and learning and motor deficits, they retain basic locomotor function, and are normally motivated to drink and eat on their own, with normal viability (Darvas, Henschen, and Palmiter 2014; Garrett Morgan et al. 2015; Henschen, Palmiter, and Darvas 2013).

We use this model, referred to as DAT:TH KO, to investigate transcriptome changes in response to DA loss. Exquisite mechanisms maintain DA tone during normal physiology, with regulation at nearly all steps of DA synthesis, release and reception of the DA signal. These steps include regulation of synthesis and feedback inhibition of the rate limiting DA biosynthetic enzyme TH, regulation of size and content of synaptic vesicles, regulation of DA reuptake via DAT, regulation of D2-like DA presynaptic autoreceptors, and postsynaptic DA receptors. In addition, there can be direct or indirect effects of classical transmitters that act to modulate DA neurons (reviewed in Sulzer et al., 2016). With this degree of regulation in normal DA physiology, it is not surprising that in pathological conditions that cause loss of DA neurons, brain physiology is altered in a compensatory manner. Both in Parkinson’s Disease and in Parkinson’s Disease models resulting from agents leading to death of DA neurons, motor symptoms occur only after a majority of DA neurons are lost (Ballanger et al. 2016; Golden et al. 2013).

This compensation for DA neuron loss is not well understood (Bezard and Gross 1998; Erwan Bezard et al. 2001; Erwan Bezard, Gross, and Brotchie 2003; Blesa et al. 2012; Hornykiewicz 1975; Perez et al. 2008). Although many of these reports indicate compensatory modulation of the DA system, other transmitter systems may also be involved, including striatal preproenkephalin (Erwan Bezard et al. 2001), altered cortical-basal ganglia-thalamocortical circuitry (Erwan Bezard, Gross, and Brotchie 2003; Erica J. Melief et al. 2018), alterations of the serotonergic system, in addition to altered DA signaling and altered extracortical circuits (Ballanger et al. 2016).

Here we analyze transcriptome responses to DA loss in pre- and postsynaptic brain regions, using DAT:CRE driven inactivation of *Th*. We find a significant difference in responses to DA loss between SNc and VTA and a surprising diversity of responses between these regions. We present evidence for a DA dependent positive feedback loop in the SNc that may aid in maintenance of DA neuronal identity.

## Materials and methods

### Animals

All animal use procedures were approved by the Institutional Animal Care and Use Committee at the University of Washington. Mice were maintained on a C57Bl/6 genetic background a and were group housed under a 12-h, light–dark cycle in a temperature-controlled room with food and water available ad libitum. Mice with inactivation of the Th gene in DAT-expressing neurons (DAT:TH KO) were generated by crossing mice with two floxed *Th* alleles (*Th^lox/lox^*) with mice that have one deleted *Th* allele (Th ^/+^) and one allele with Cre recombinase expressed under the control of the *Slc6a3* gene encoding DAT (*Slc6a3^Cre/+^*). DAT:TH KO mice had the genotype *Th ^/lox^-Slc6a3^Cre/+^*. C57Bl/6 were used as wild type (WT) control animals.

### RNA Extraction

Mice, 3-5-month-old, were sacrificed by decapitation after brief exposure to CO2; the VS, DS, SNc, VTA and PFC from each mouse were dissected from the brain. We collected standardized tissue punches from projection fields (1 mm diameter) using the mouse brain atlas and a brain-matrix to generate standardized slices (2 mm thick for striatum and 1 mm thick for PFC). For the midbrain, we collected slices similar to those described for the projection-field punches and then dissected SNc and VTA similar to what is described in Lautenschläger et al. (2018). Tissues were immediately frozen in liquid nitrogen and stored at −80 °C. Tissues were processed, and subsequently cDNA libraries from isolated RNA were prepared in three independent batches; first one with samples from 3 DAT:TH KO and 3 WT animals; second with samples from one mouse per group; third with 2 DAT:TH KO and 2 WT animals. Each time tissues were homogenized using Bullet Blender with 0.5 mm RNase Free Zirconium Oxide Beads (Next Advance) in RLT buffer from RNeasy Plus Micro Kit (Qiagen). Total RNA was isolated using the RNeasy Plus Micro Kit (Qiagen). The quality and quantity of RNA from each sample was determined using an Agilent 2100 Bioanalyzer with Agilent RNA 6000 Pico Chips. Messenger RNA was then obtained from at least 100 ng of total RNA using NEBNext® Poly(A) mRNA Magnetic Isolation Module. RNA-seq libraries were prepared using NEBNext® Ultra™ Directional RNA Library Prep Kit for Illumina® following manufacturer’s protocol. The library quality and insert length were checked using the Agilent 2100 Bioanalyzer with DNA High Sensitivity DNA Kit. The first two sets of 30 and 10 libraries were sequenced using Illumina HiSeq 2000 (BGI), third set of 20 libraries was sequenced using Illumina HiSeq 2500 (UCLA).

The quality of obtained fastq files was examined using FASTQC (S. 2010). The reads were aligned to the mouse reference genome (GRCm38) using HISAT2 v2.1.0 (D. Kim, Langmead, and Salzberg 2015) aligner with --rna-strandness RF parameter. Aligned reads were then summarized with featureCounts from the SourceForge Subread package (Liao, Smyth, and Shi 2014). Summarizing, we obtained over 2.3 billion uniquely mapped 100 bp paired-end reads (42206643± 4754672 uniquely mapped reads per library). Sequencing data have been deposited in the Sequence Read Archive under accession code PRJNA550435.

### Statistical analysis

According to the current bioinformatic standards required for performing statistically satisfying RNA-Seq experiment detecting low abundance differentially expressed transcripts at p<0.05 we needed 3 biological replicates of each genotype. However, due to unexpected phenotype of half of the DAT:TH KO samples, we increase the sample size to 6 biological replicates of each genotype. A differential expression analysis was performed with R/Bioconductor package DESeq2. The negative binomial distribution, with the batch variable included into the model design, was fitted to raw counts output from featureCount (Love, Huber, and Anders 2014). Following, the Wald test was used for significance testing. Multiple testing was corrected for using the Benjamini-Hochberg procedure. The hierarchical cluster analysis was preformed using Euclidean distance metrics.

## Results

### A novel *Th* partial transcript can be detected in DAT:TH KO presynaptic libraries

We used the DAT:TH KO model to test for effects of striatal DA deficiency on the transcriptome in the regions studied. We expected that the lack of DA will affect preferentially DA producing and DA receiving parts of the brain. Therefore, we studied transcripts from five distinct brain regions, two of which are presynaptic: *Substantia nigra pars Compacta* (SNc) and Ventral tegmental area (VTA); and three postsynaptic: dorsal striatum (DS), ventral striatum (VS) and prefrontal cortex (PFC), from six wild type and six DAT:TH KO mice. Samples from each brain region were sequenced with similar depth of 42.2 ± 4.75 million uniquely mapped reads per library. The distributions of number of reads mapped per gene in all brain regions followed the same pattern, with the majority of genes represented by over 1000 unique reads (Fig. S1A). In all the following analyses we filtered out weakly expressed genes as represented by fewer than 30 reads total in all samples from a given brain region. In the final dataset, a majority of genes (14,157) was detectable in the samples from all brain regions, but there are 33 to 142 genes that are found in only one region (Fig. S1B).

To validate an effective *Th* knockout in DAT expressing neurons of the DAT:TH KO animals, we examined the expression level of *Th* transcripts in the SNc and VTA samples. Unexpectedly, we detected a high level of *Th* mRNA in SNc and VTA from three DAT:TH KO mice which we designate as ‘DAT:TH KO^High^*’* versus ‘DAT:TH KO^Low^*’* (Fig. 1A). Additionally, SNc and VTA samples from one WT animal have remarkably low levels of *Th* expression. To determine if this low level of *Th* mRNA correlates with the DAT:TH KO^Low^ samples, we mark samples from this WT animal with minus sign (“-”) in this and subsequent figures.

**Figure 1.**
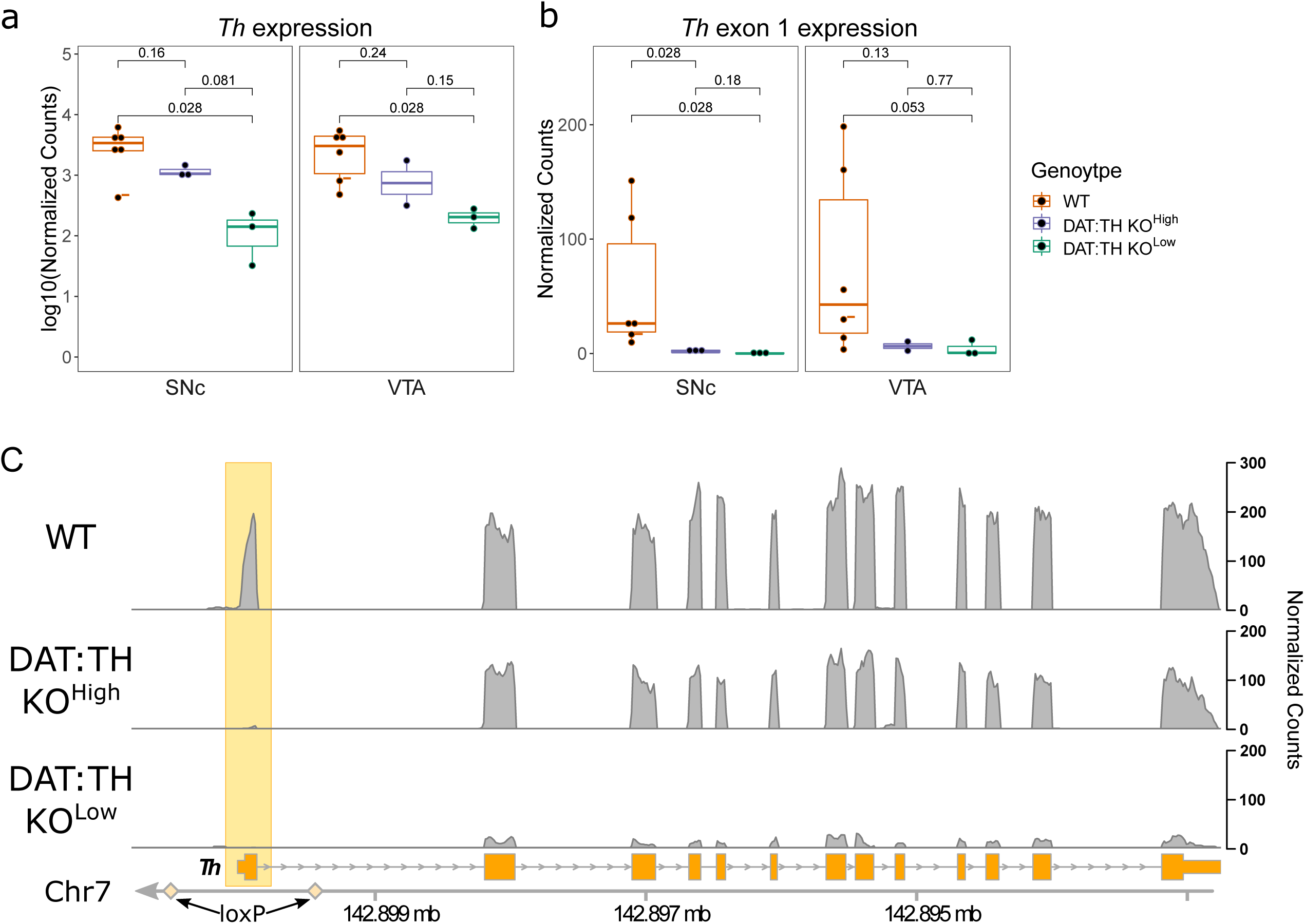
Th expression level in SNc and VTA samples. DAT:TH KO animals were split into two groups: DAT:TH KO^High^ or DAT:TH KO^Low^. DAT:TH KO animals were assigned to DAT:TH KO^High^ group if they had *Th* expression level higher than any WT animal in a given brain region. In all following analyses all samples from the same animal are grouped into the same DAT:TH KO^Low or High^ group. DESeq2 normalized counts are based on featureCounts output. A) Normalized expression level of Th in both presynaptic regions. The normalized counts are shown on log10 scale. B) Normalized expression level of Th exon 1 is significantly lower in all DAT:TH KO animals in comparison to WT. Numbers over bars represent p-value from Wilcoxon tests. Datapoints from “WT-” mouse are marked with “-”. C) Representative per base coverage plot of Th. The bottom part of the plots shows known Th exons in orange and yellow rhombuses mark loxP sites flanking exon 1. Yellow square marks the first exon, that should be removed by DAT:Cre recombinase in DAT:TH KO animals. Only few Th exon 1 reads are detectable in all DAT:TH KO samples, however the remaining exons have a high level of coverage in the DAT:TH KO^High^ SNc and VTA samples.

One could imagine that inefficient removal of the *Th* exon 1 by Cre recombinase by low level of Cre expression, could explain the observed variation in levels of TH transcripts over the DAT:TH KO samples. However, we detect very few exon 1 reads in all of the DAT:TH KO samples (Fig. 1B), even in the DAT:TH KO samples with high *Th* expression level (DAT:TH KO^High^) which are otherwise of normal downstream structure (Fig. 1C).

We used two analytic methods, PCA and hierarchical cluster analysis, to determine whether the SNc transcriptomes of WT, DAT:TH KO^High^ and DAT:TH KO^Low^ show divergence in their overall transcriptomes. The PCA and the hierarchical cluster analysis from SNc show that DAT:TH KO^High^ have overall expression pattern more similar to wild type than to the DAT:TH KO samples with low level of *Th*. The WT and DAT:TH KO^High^ samples from SNc are localized closer to the upper-left corner of the PC1 vs PC2 plot, whereas DAT:TH KO samples are grouped near lower-right corner (Fig. 2A, a cluster of samples with low level of *Th* is outlined with dashed line). This separation is even more visible on the plot summarizing the hierarchical cluster analysis (Fig 2B). DAT:TH KO samples with low level of *Th* creates one cluster with the aforementioned WT sample with the lowest level of Th (“WT-”), whereas DAT:TH KO samples with high level of Th are grouped with the remainder of the WT samples on the other branch of dendrogram.

**Figure 2.**
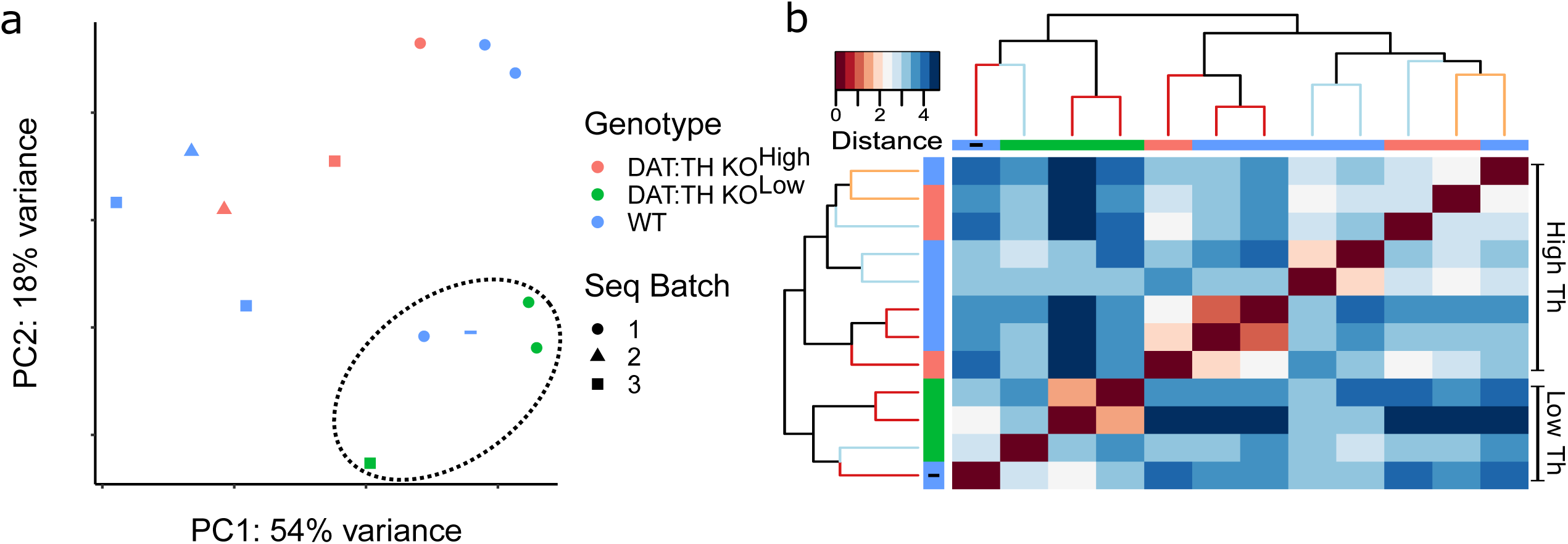
Clustering analyses of SNc samples. A) SNc PCA based on genes represented by at least 30 normalized counts across all samples. The datapoint from “WT-” mouse is marked with “-”. The dashed ellipse outlines a cluster of samples with low level of *Th*. B) Hierarchical cluster analysis of SNc samples. Dendrogram and heatmap were created based on the maximum distance between samples. Blue color indicates large distance, with red diagonal line representing identity from comparison of the same sample. Each row and column represent an individual library, with row/column labeling by color coded boxes, following the same pattern as PCA plot. Leaves of dendrogram are color coded by sequencing batch. Blue squares with “-” mark the sample form “WT-” mouse.

### Lack of striatal dopamine does not alter the overall pattern of gene expression

Upon finding variation between the DAT:TH KO samples, we examined general differences between brain regions and genotypes by two techniques, principal component analysis (PCA) (Fig. 3A) and hierarchical cluster analysis (Fig. 3B). The first principal component in PCA explains over the half of the variability, and clearly separates pre- and postsynaptic regions. The second principal component separates SNc, VTA, DS and VS from PFC. On the contrary, WT vs DAT:TH KO genotype is not recognizable from this analysis, indicating that the effect size of the DA deficiency is smaller than the effect size of brain region. All samples from both presynaptic regions, SNc and VTA, are closely localized, independent of genotype, reflecting similarity in their gene expression profiles. The libraries from DS and VS create another sparse group, and the PFC samples form an independent group, reflecting their separate origin. However, one sample from DAT:TH KO PFC (Fig. 3A, datapoint indicated by a hollow square) did not localize closely to the PFC group. The library used to generate this datapoint had only 20% of the expected read number, with 30% of reads representing poly-G, suggesting a problem with library preparation or sequencing. None of the other libraries showed these issues, so this library was eliminated from further consideration. Similar strong grouping by a region of origin is visible in a heatmap representing results of hierarchical cluster analysis (Fig. 3B). Two main branches of the dendrogram separate pre- and postsynaptic samples, with a further division into samples from PFC and striatum. As with the PCA, the DAT:TH KO and WT samples do not show clustering. Moreover, we do not detect any strong batch affect and samples from three different sequencing batches are clustered in alternating order (color coded in the leaves of the dendrogram in Fig. 3B).

**Figure 3.**
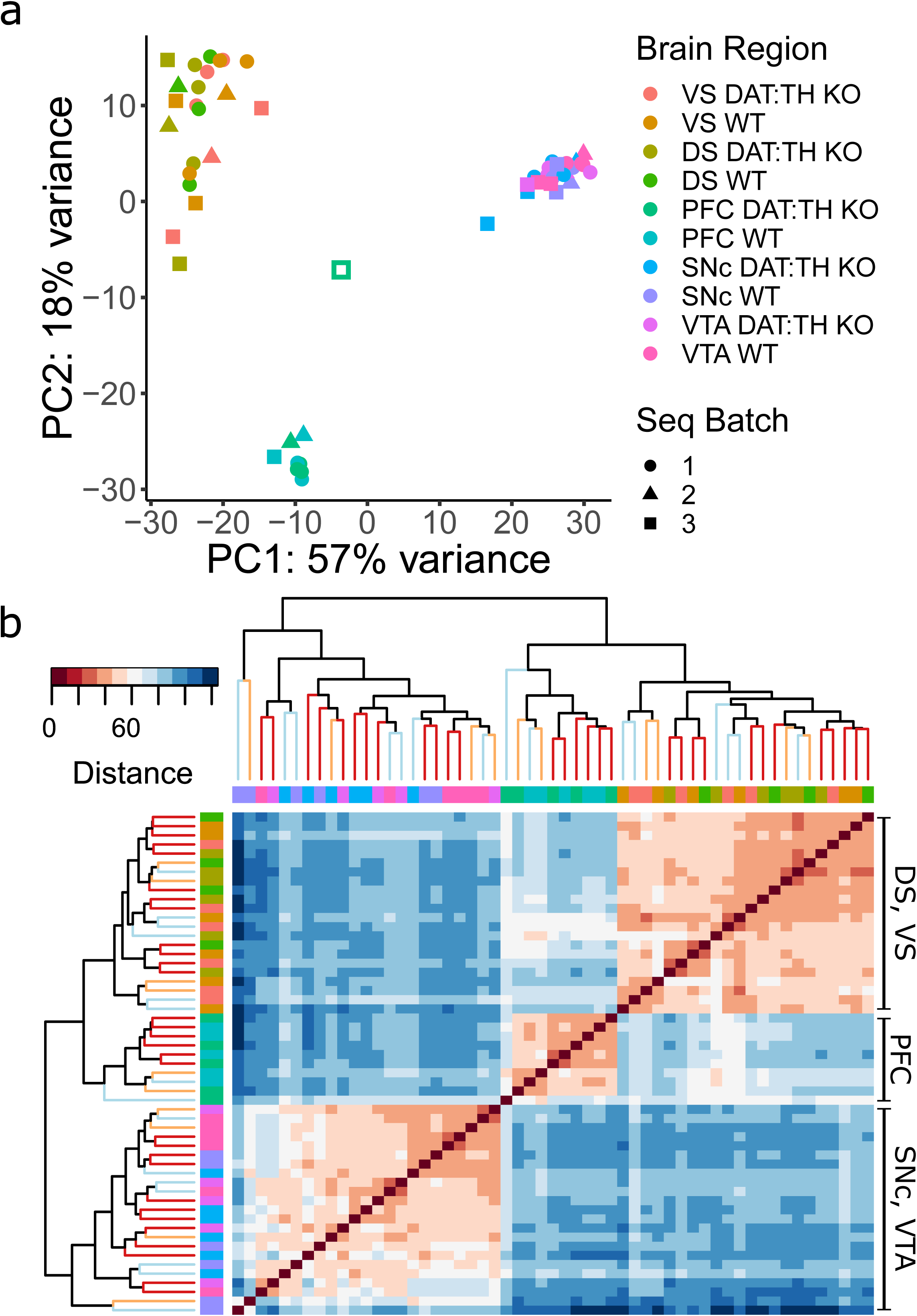
Cluster analyses. Input data for all analyses are DESeq2 normalized count values based on featureCounts output. A) Principal Component Analysis of all sequenced samples from three independent batches of RNA-sequencing, coded by the shape of datapoints. Each data point represents an individual library, color coded based on the brain region and genotype. The hollow square marks one outlier PFC library. B) Hierarchical cluster analysis of all sequenced samples from three independent batches of RNA-sequencing. Dendrogram and heatmap were created based on the Euclidean distance between samples. On heatmap blue color indicate large distance, with red diagonal line representing identity from comparison of the same sample. Each row and column represent an individual library, with row/column labeled by color coded boxes based on the brain region, following the same pattern as PCA plot. Leaves of dendrogram are color coded by sequencing batch: 1^st^ – red, 2^nd^ – yellow, 3^rd^ – light blue.

### Lack of DA results in small transcriptomic effects in postsynaptic brain regions

To address whether DA deficiency affects gene expression in the DS, VS and PFC, we compared transcriptomes from the DAT:TH KO^Low^ and WT animals using a DESeq2 statistical analysis based on a negative binomial distribution model. We analyzed the expression level of 14,648, 14,694 and 14,692 genes per brain region, respectively. DS samples have the strongest observed response to the decreased DA level among the postsynaptic brain parts, with 7 differentially expressed (DE) genes at a false discovery rate (FDR) < 0.05. We detected only 3 downregulated: *Nefm*, *Pdyn* and *Sntg2* (Fig. 4A); and only 4 upregulated genes in DS: *Anxa3*, *Fam43a*, *Slc7a8*, and *Wdfy* (Fig. 4B). Only one of those genes, *Sntg2*, is also differentially expressed in the ventral striatum of DAT:TH KO^Low^ animals. Additionally, *Gpx6* has a significantly higher level of expression and *Shisa8* is significantly downregulated in DAT:TH KO^Low^ VS samples. Finally, we do not detect any DE genes in DAT:TH KO PFC samples.

**Figure 4.**
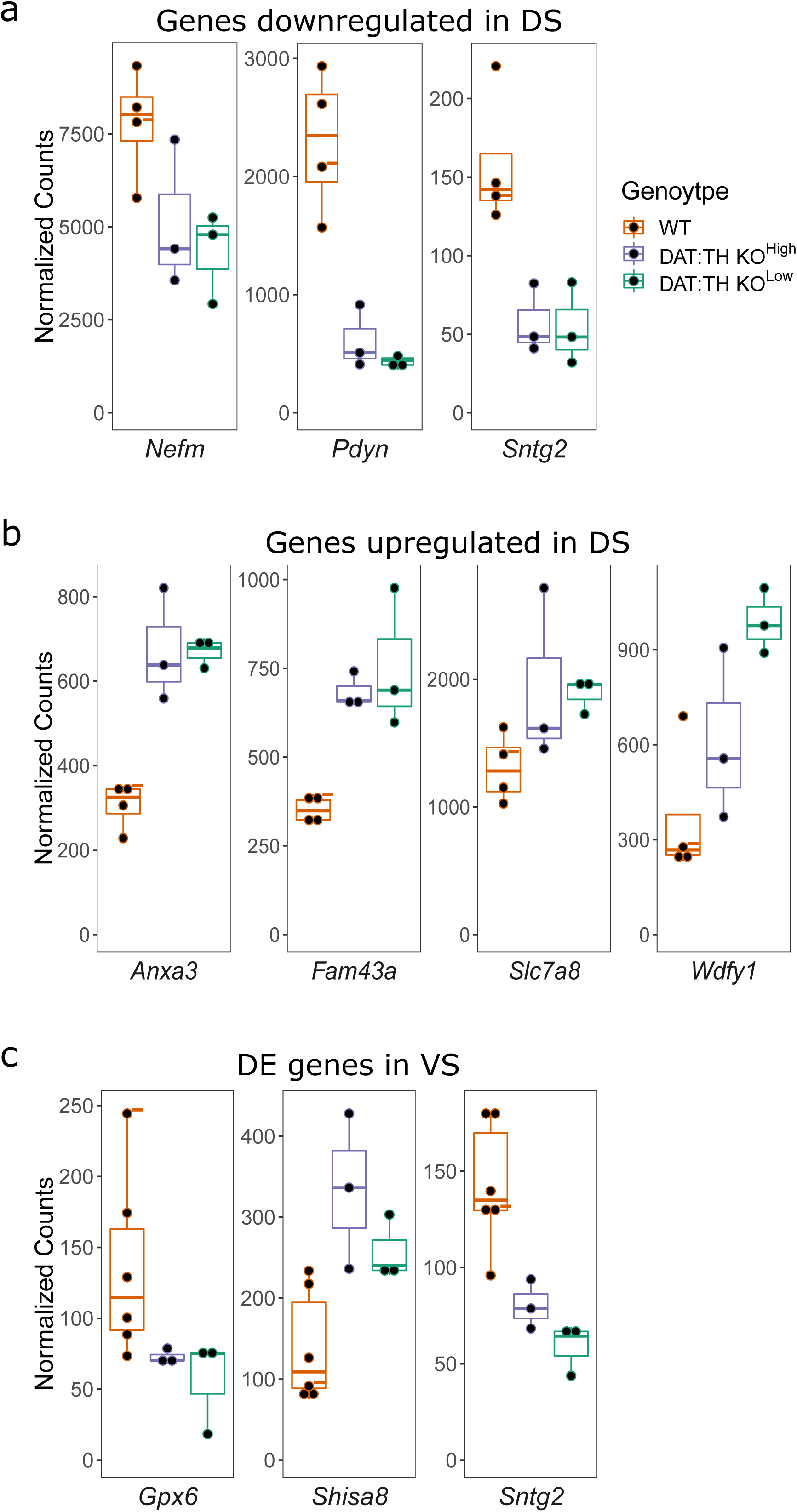
Expression level of DE genes in postsynaptic brain regions. All plotted genes have statistically significant DE in DAT:TH KO^Low^ at FDR <0.05. All figures follow color coding from figure A. Datapoints from WT mouse with low level of *Th* mRNA are marked by “-” sign. The lower number of WT samples in DS is due to a loss of 2 libraries during preparation. A) Genes downregulated in DS. B) Genes upregulated in DS. C) DE genes in VS.

### DAT:TH KO^Low^ show reduced expression of DA related genes in SNc

We next asked whether DA deficiency leads to differential gene expression in the SNc. Comparing expression level of 15,282 genes from the DAT:TH KO^Low^ and WT animals, we find 34 that are DE at a false discovery rate (FDR) < 0.05 (Fig. 5A). The DAT:TH KO^High^ libraries were not used during the detection of the DE genes since earlier analyses show that these libraries most resemble WT. However, when we examined the expression level of the DE genes from the DAT:TH KO^High^ with the DAT:TH KO^Low^ and the WT SNc libraries, DAT:TH KO^High^ cluster with the WT samples (Fig. 5B), indicating that TH expression level is a better predictor of clustering than genotype. When we directly compared the DAT:TH KO^High^ and the WT SNc samples, none of the previously detected downregulated genes except for *Lcn2* and *Lrg1* showed significant DE.

**Figure 5.**
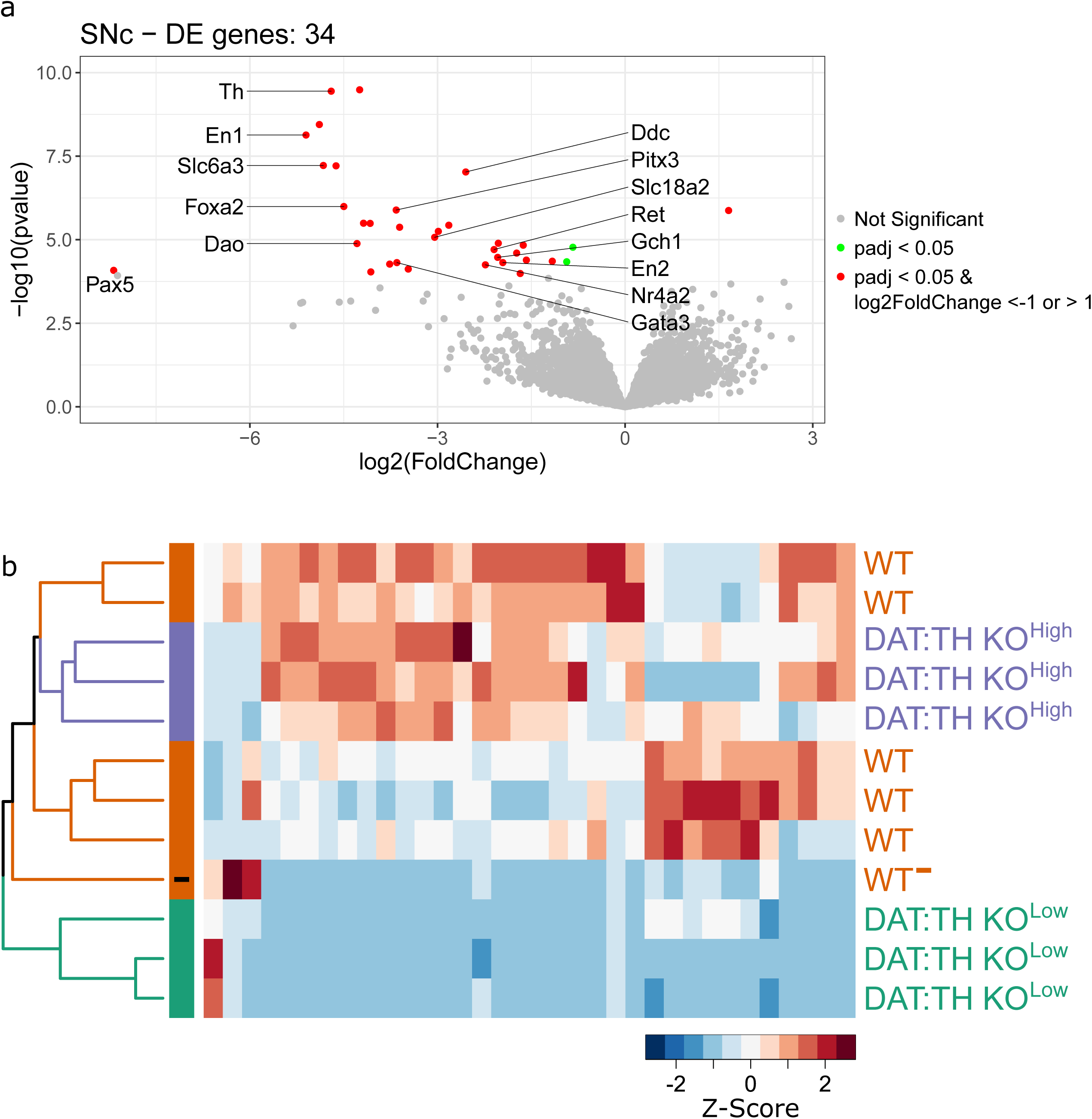
DE genes in SNc. A) Volcano plots of DE analysis between WT and DAT:TH KO^Low^ samples. Data points are color coded based on their adjusted p-value at FDR <0.05 and fold change. DE DA-related genes are labeled. B) Heatmap representing hierarchical clustering of the SNc libraries based on detected DE genes. Each column represents an expression patter of one DE gene. Color scale represent Z-score, with red marking high expression level. Leaves of dendrogram and row labels are by color coded based on the genotype and expression level of Th. WT sample with low level of *Th* mRNA is marked by “-” sign.

All but one of the DE genes are downregulated in the DAT:TH KO^Low^ SNc samples (Fig. 5A). The upregulated gene, 2900052N01Rik, encodes a long intergenic noncoding RNA with no known function. For most genes we observed a ∼10-fold expression level decrease (with median log2 FoldChange of −3.47 and mean log2 FoldChange of −3.2) in the DA depleted SNc. Among the downregulated genes are several transcription factors involved in DA neuron specification: *Foxa2*, *Gata3*, *Nr4a2*, *Pax5*, *Pitx3*, *En1* and *En2* (Fig. 6A). Consistent with decreased expression of the crucial dopaminergic transcriptional regulators, there is a reduced level of classical dopaminergic markers: *Dao*; *Ddc*; *Ret*; *Gch1*; *Slc6a3* (DAT); *Slc18a2* (VMAT2) (representative expression plots are collected on Fig. 5B). Supplemental file 1: Table S1 reports the complete results of this analysis, providing full information about all DE genes.

**Figure 6.**
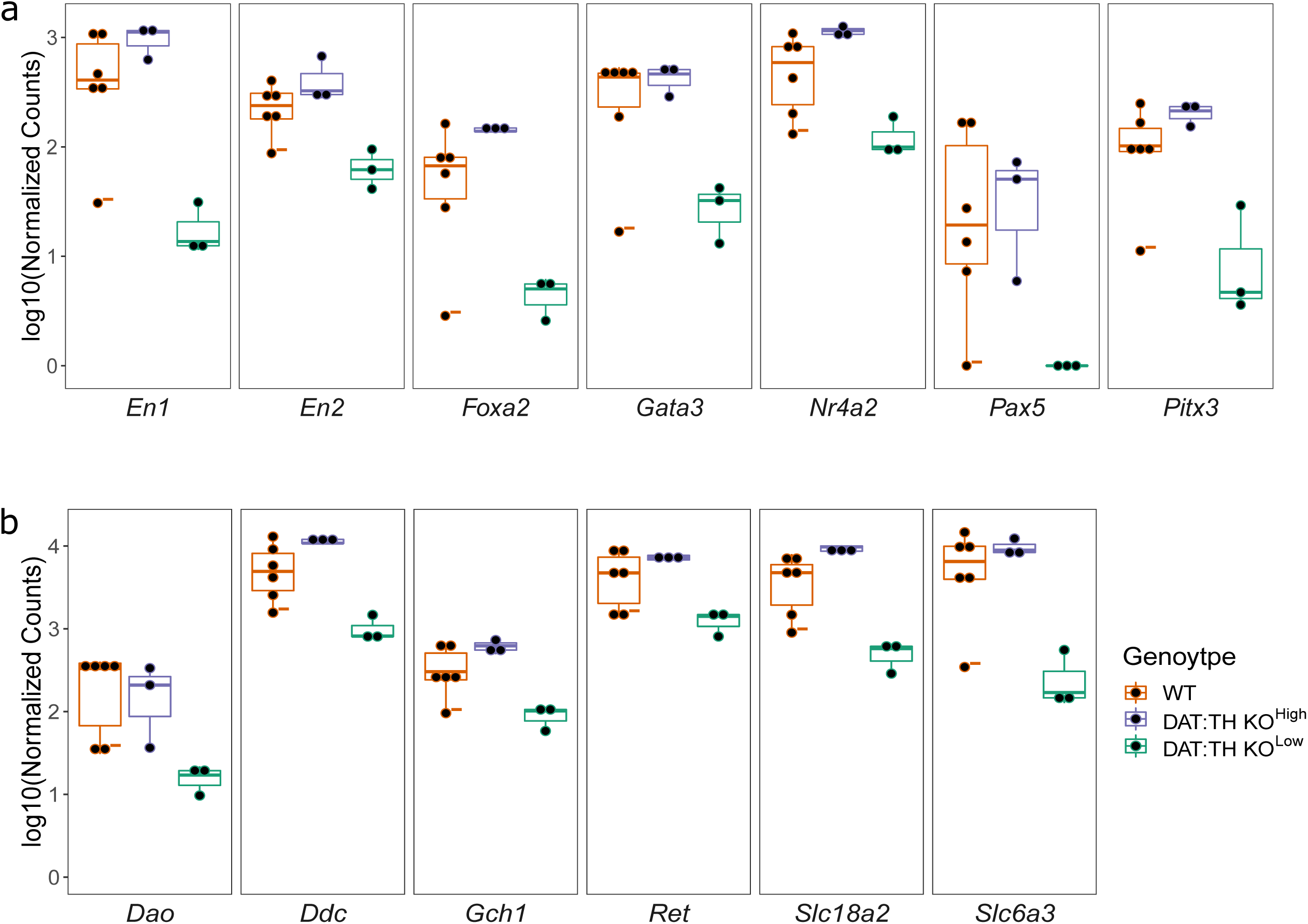
Representative expression level of genes DE in SNc. All plotted genes have statistically significant decreased expression level in DAT:TH KO^Low^ vs WT samples at FDR <0.05. All figures follow color coding from figure B. Datapoints from WT mouse with low level of *Th* mRNA are marked by “-” sign. A) log_10_(Normalized Counts) of DA-related downregulated transcription factors. B) log_10_(Normalized Counts) of representative downregulated dopaminergic markers.

### VTA and SNc Transcriptomes Respond Distinctly to DA Deficiency

Comparing expression levels of 15,187 genes detected in VTA in the DAT:TH KO^Low^ and WT samples, we detected 89 DE genes at FDR < 0.05. In contrast to the general directionality of the DE observed in SNc, the majority of DE genes were upregulated in DAT:TH KO^Low^ VTA samples (55 upregulated vs 34 downregulated genes; Fig. 7A). Moreover, the magnitude of observed changes is smaller than in SNc. The majority of the DE is down- or upregulated less than two-fold (60 out of 89 DE genes, with median absolute value of log2 Fold Change of 0.7 and mean of 1.09). However, the hierarchical cluster analysis based on the DE genes shows again that all DAT:TH KO^Low^ samples are clustered together, whereas DAT:TH KO^High^ are more similar to WT. Moreover, the “WT-” sample with a relatively low level of the *Th* expression is clustered with the DAT:TH KO^Low^ libraries, showing an intermediate pattern of DE (Fig. 7B). Furthermore, when we compared DE across the entire transcriptome, none of the DE genes detected between DAT:TH KO^Low^ and all WT samples have a statistically significant difference in expression level between DAT:TH KO^High^ and WT VTA.

**Figure 7.**
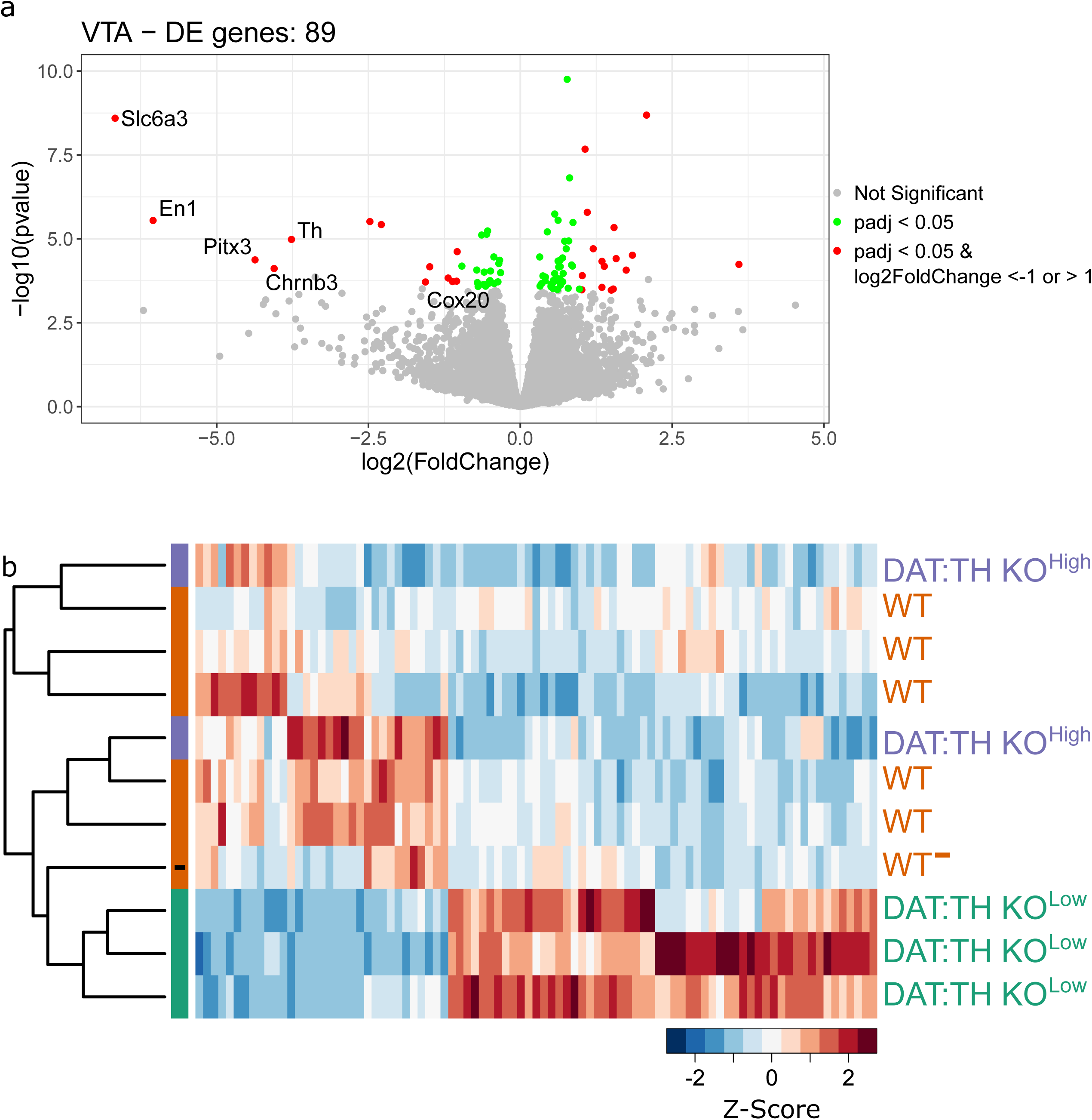
Genes DE in VTA. A) Volcano plots of DE analysis between WT and DAT:TH KO^Low^ VTA samples. Data points are color coded based on their adjusted p-value at FDR <0.05 and fold change threshold. DE DA-related genes are labeled. B) Heatmap representing hierarchical clustering of the VTA samples based on detected DE genes. Each column represents an expression patter of one DE gene. Color scale represent Z-score, with red marking high expression level. Leaves of dendrogram and row labels are by color coded based on the genotype and expression level of *Th*. WT sample with low level of *Th* mRNA is marked by “-” sign.

In addition to *Th*, only five DE genes from DAT:TH KO^Low^ SNc were also significantly downregulated in VTA samples: *Chrnb3*, *Cox20*, *En1*, *Pitx3* and *Slc6a3* (Fig. 8A). Among the remaining downregulated genes in the VTA, only two are known transcription factors: *Foxa1* and *Pbx3*. Thus, no additional classical dopaminergic markers were downregulated in VTA.

**Figure 8.**
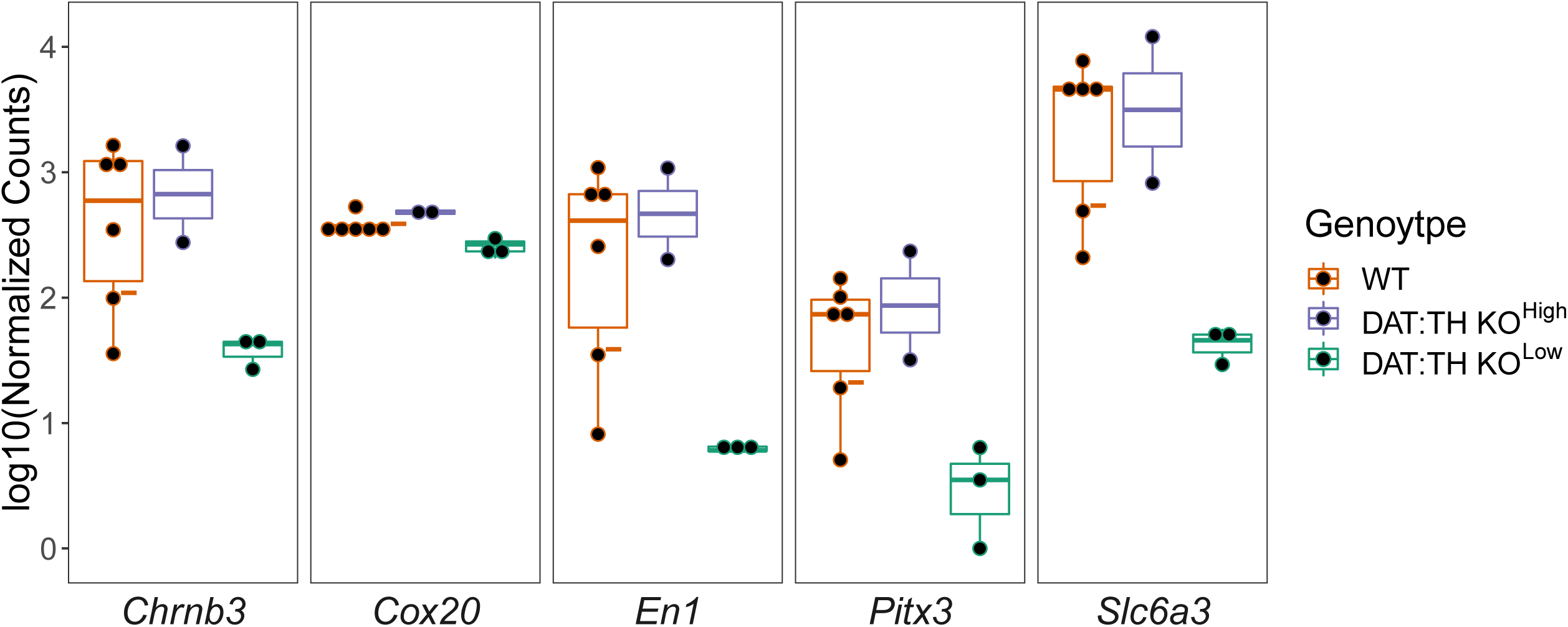
Expression levels of genes downregulated in both DAT:TH KO presynaptic regions across VTA samples. All plotted genes have statistically significant decreased expression level in DAT:TH KO^Low^ SNc and VTA at FDR <5%. Datapoints from WT mouse with low level of *Th* mRNA are marked by “-” sign.

To further examine potential differences between the dopamine dependency in SNc and VTA we looked at the Gene Ontology (GO) terms enriched in detected DE genes, with top enriched Biological Process (BP) GO terms are collected in Table 1. The full results of the GO enrichment analyses are shown in the supplemental material (Additional file 2, Table S2). As expected, the top ten most enriched terms in genes downregulated in SNc are connected to DA metabolism and to dopaminergic neurons and their functions (20 out of 34 enriched categories). Results of the enrichment analysis from the genes downregulated in the VTA are not as explicit. Only 5 categories were significantly enriched after correction using the Benjamini–Hochberg procedure. While none of them directly connect to DA, one term, functionally related to DA, the GO:0007626 (locomotory behavior), is also enriched in DE genes in SNc. The other two related categories are connected to mitochondria. In contrast to the downregulated, the genes upregulated in VTA show high level of enrichment in 85 BP categories. This difference could stem from the greater number of genes with increased level of expression in DAT:TH KO^Low^ VTA samples. However, it could also suggest stronger and active upregulation of particular pathways. The top ten enriched terms revolve around synaptic transmission and glutamatergic signaling. Other terms of particular interest point to the regulation of calcium and GABA-ergic signaling. These results suggest that while the decrease level of DA in SNc may cause loss of dopaminergic identity, the neurons in VTA seem to have more synaptic plasticity and react by remodeling of synaptic signaling.

**Table 1.**
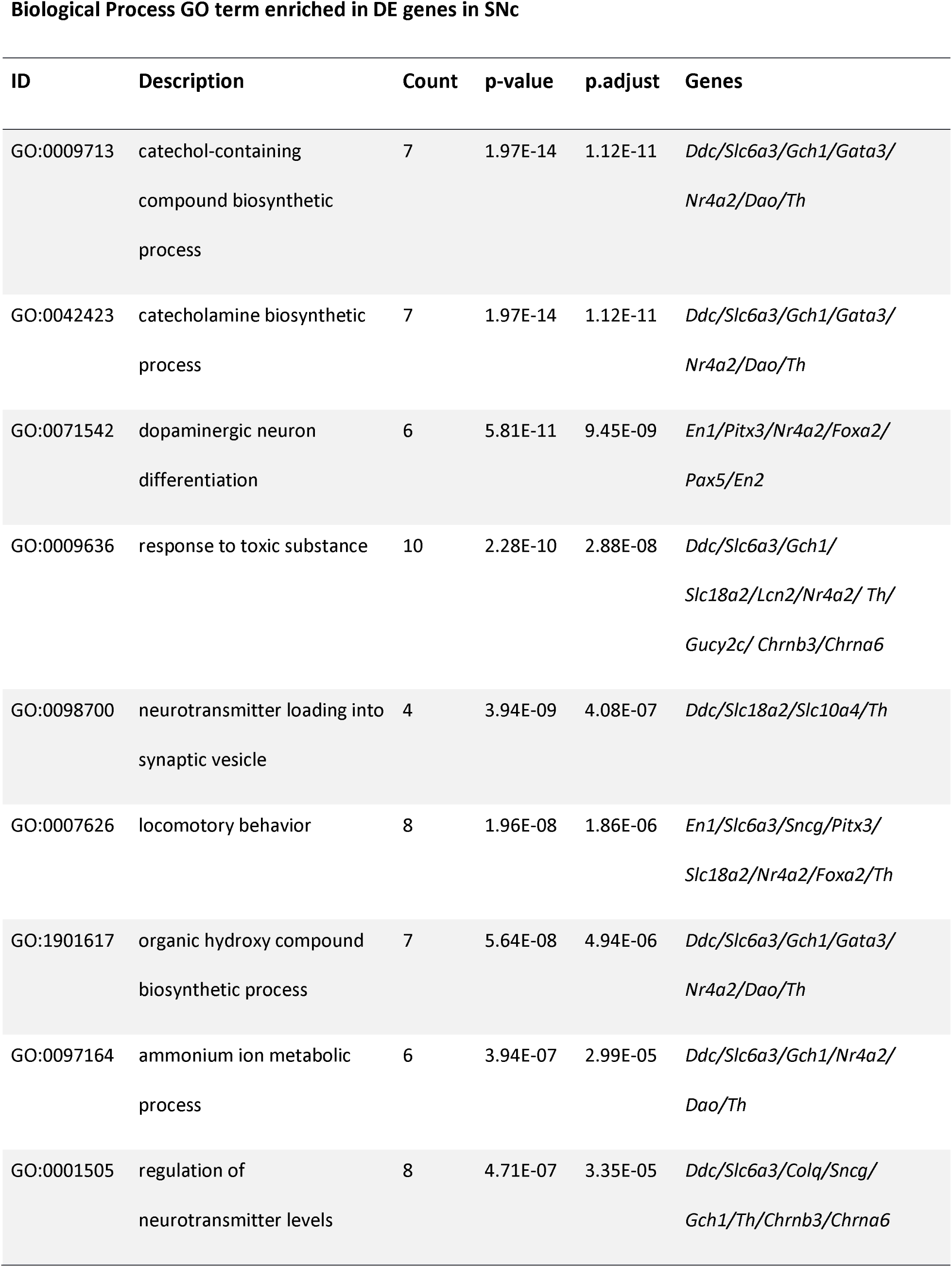
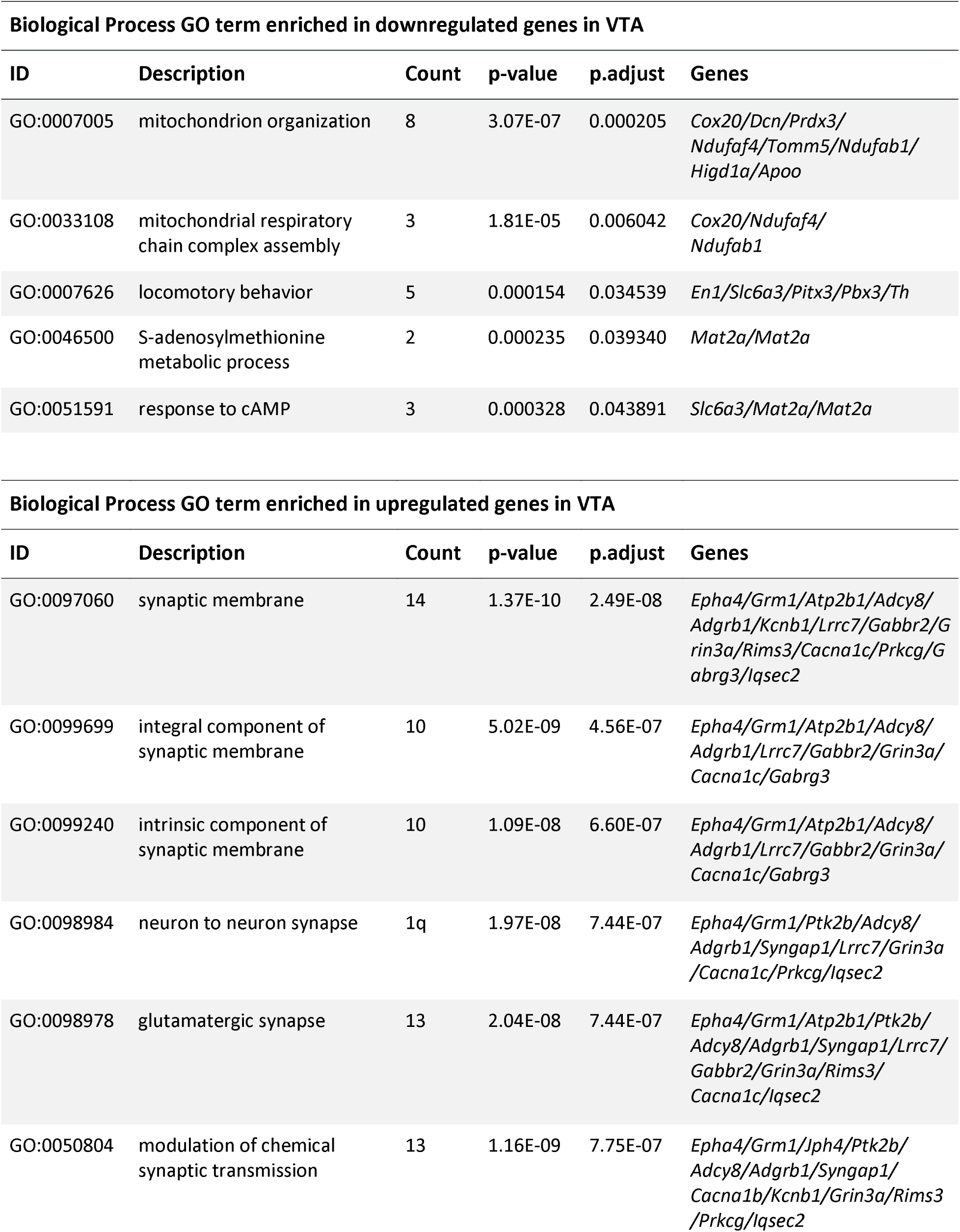
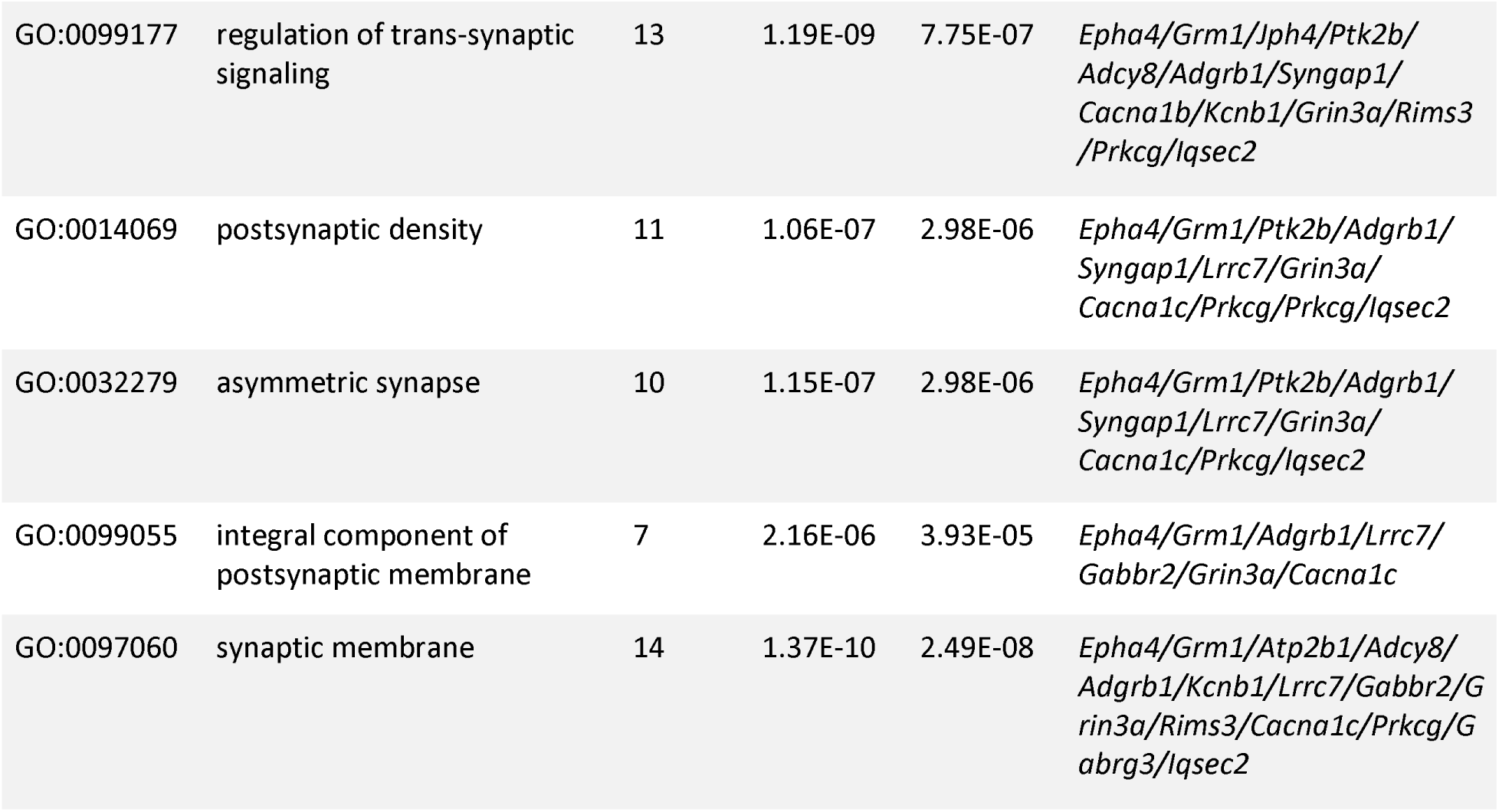
Results of the GO enrichment analysis based on the DE genes in SNc and VTA at FDR < 0.05. P-adjusted values were corrected using Benjamini–Hochberg procedure. This table shows only up to the top ten terms enriched based on the DE gene in SNc and VTA, from the total number of 33 and 85 significantly enriched categories.

## Discussion

In this study we used a DAT:TH KO mouse model, generated by DA transporter driven CRE to inactivate the *Th* gene, with loxP sites flanking *Th* exon 1, which encodes the *Th* translation startsite. This approach results in animals that lack 95% of normal levels of DA in the striatum, whereas the number of DA neurons is otherwise unaffected (Darvas, Henschen, and Palmiter 2014; Garrett Morgan et al. 2015). This allows observation of transcriptome changes in response to DA loss, especially in SNc and VTA, that could have been missed in models based on the loss of DA neurons. Furthermore, as the DAT:TH KO model used here does not induce neuronal death, we could separate effects of the DA deficiency and a decreased amount of DA projections into striatum that are difficult to discriminate in the lesion-based models. Previously published comparisons between these two approaches show that more severe symptoms of PD, such as bradyphrenia, cognitive rigidity and spatial learning deficits, are dependent on both loss of DA and DA neuronal loss in striatum (Darvas, Henschen, and Palmiter 2014). However, cognitive flexibility, working memory and cue-dependent associative learning can be caused solely by a decrease in striatal DA, independent of depletion of nigrostriatal DA neurons (Garrett Morgan et al. 2015).

Surprisingly, we detected near normal levels of *Th* transcripts in half of the DAT:TH KO samples (Fig. 1A). However, almost no reads were mapped to the FLOXed *Th* exon 1 in either of the DAT:TH KO samples, eliminating the possibility that these unexpectedly high levels of Th transcripts result from inefficient Cre recombinase function (Fig. 1B and C). The few *Th* exon 1 reads in DAT:TH KO samples could derive from DA neurons that do not express DAT (Darvas, Henschen, and Palmiter 2014).

The unusual *Th* transcript that is detectable in the DAT:TH KO^High^ samples lacks the first exon that encodes the normal translation startsite, yet the remaining transcript has the same structure as native *Th* mRNA. The normal N-terminal domain, encoded by exon 1, is involved in regulation of TH enzymatic activity, containing several phosphorylatable serines involved in TH activation (Tekin et al. 2014), though deletion of this region can stabilize the TH protein (Nakashima et al. 2005). To date, only one isoform of *Th* was characterized in mouse, however, there are four alternative splicing forms of *TH* in humans that differ in structure at the 5’ end of the transcript, resulting in proteins with different N-terminal regulatory segments (Tekin et al. 2014). The *Th* transcripts detected in DAT:TH KO^High^ animals could potentially represent a novel mouse *Th* isoform that is differentially regulated relative to wildtype *Th*. The regional PCA and hierarchical cluster analyses from SNc samples show that DAT:TH KO^High^ cluster more strongly with WTs than with DAT:TH KO^Low^ animals (Fig. 2), supporting this hypothesis. Therefore, we decided to treat the samples from those animals as a separate group, that could represent a potential DA-compensatory mechanism. Interestingly, when we compared the expression levels of genes differentially expressed between DAT:TH KO^Low^ and WT, the samples from DAT:TH KO^High^ resemble WT animals. In the hierarchical cluster analysis (Fig. 5 and 7) all DAT:TH KO^High^ samples clustered between WT samples. Additionally, WT samples with lower level of Th, and thus presumably lower level of DA, are clustered closer to the DAT:TH KO^Low^ animals. These results support our hypothesis that the alternative Th transcripts from DAT:TH KO^High^ animals could rescue the DAT:TH KO^Low^ phenotype. However, further studies will be needed to confirm if DAT:TH KO^High^ animals exhibit more wildtype-like behavioral phenotypes and whether they express normal levels of DA.

In our study we detected nine genes DE in the postsynaptic regions, seven in DS and three in VS, with one gene downregulated in both parts of striatum. These findings suggest that the previously described broad transcriptional changes in striatum in neurodegeneration-based PD models result from decreased numbers of DA neurons, and not only by the decrease in DA level (Chin et al. 2008; Patel et al. 2008; Pattarini et al. 2008; Singh et al. 2010). We confirmed the dopaminergic regulation of dynorphin expression in DS described previously in DA deficient models based on both complete inactivation of *Th* and 6-OHDA-lesions (Sivam 1996; Zhou and Palmiter 1995). Dynorphin belongs to an endogenous opioid peptide family, involved in regulation of DA release, motor behavior, and learning (Carey et al. 2009; Gerfen and Steiner 1998; Reid et al. 1990; You, Herrera-marschitz, and Terenius 1999). The transcriptional activity of *Pdyn*, the gene encoding prodynorphin, is dependent on signaling through the DA D1 receptor, and works in a regulatory loop suppressing excessive activation of the striatal neurons (Gerfen and Steiner 1998; M. Xu et al. 1994). We detected decreased expression of a related protein, syntrophin gamma 2, encoded by *Sntg2*, in both DS and VS DAT:TH KO^Low^ samples. Syntrophin gamma 2 belongs to the family of cytoplasmic peripheral membrane proteins that binds to the C-terminal of dynorphin and act as adaptor proteins by its PDZ domain (Alessi et al. 2006; Piluso et al. 2000). However, target proteins associated with dynorphin-syntrophin scaffold complex are currently unknown. Surprisingly, the striatal DAT:TH KO^High^ samples have comparable levels of expression of DE genes as DAT:TH KO^Low^ samples. We suspect that this similar response is due to all TH KO transcripts lacking the coding region of exon 1. This region encodes the sequence directly upstream of serine 31 whose phosphorylation targets tyrosine hydroxylase to vesicles for transport along microtubules (Jorge-Finnigan et al. 2017). Without this axonal transport, the DAT:TH KO^High^ mice would be expected to show no rescue in the striatal regions.

Looking more closely at differences between SNc from animals with low levels of *Th* mRNA relative to WT controls, we detect 33 strongly downregulated genes (Fig. 5), of which 14 are dopaminergic markers and/or genes required for DA neurons differentiation or maintenance. This result leads to the hypothesis that this decreased gene expression is due to loss of DA signaling, implying a positive role for maintaining DA neuron identity. Downregulation of the DA related genes in SNc of DAT:TH KO^Low^ animals is in accord with changes in PD and the lesion-based PD animal models, where decreased expression is an expected consequence of DA neuron loss. These genes include dopaminergic markers, such as *Aldh1a7*, *Ddc*, *En1*, *En2*, *Foxa2*, *Nr4a2* (*Nurr1*), *Pitx3*, *Slc18a2* (VMAT2), *Slc6a3* (DAT) and *Th* (Domanskyi et al. 2014; Jacobs et al. 2011; Tekin et al. 2014).

A potentially important transcription factor that is involved in the development of DA neurons, *Pax5* (Simon et al. 2003), is also significantly downregulated in DAT:TH KO^Low^ SNc. This suggests that some of the observed changes could be connected to decreased striatal DA levels during development. The downregulation of the postmitotic selector *Gata3*, involved in differentiation of SNc and VTA GABAergic neurons, indicates a potential developmental effect of DAT:TH KO (Haugas et al. 2016; Moriguchi 2006).

The results of differential gene expression analysis of DAT:TH KO^Low^ and WT VTA samples highlight the important differences between VTA and SNc. We not only detect more DE genes in the VTA, but moreover the majority of them are <u>upregulated</u> in the absence of DA. This distinct response could reflect a differential variability of DA neurons in SNc and VTA in PD (Brichta and Greengard 2014), as only four DA genes have significantly lower level in DAT:TH KO^Low^ VTA (*Th*, *Slc6a3*, *Pitx3* and *En1*; Fig. 7).

Among the genes downregulated in SNc, but not in VTA, *Nr4a2*, encoding the highly conserved transcription factor Nurr1, is of particular interest. In PD patient brains, Nurr1 is downregulated in regions affected by α-synuclein inclusions and variants of *Nr4a2* are associated with PD (Grimes et al. 2006; Le et al. 2003; P. Y. Xu et al. 2002; Zheng, Heydari, and Simon 2003). Nurr1 is expressed in the early postmitotic neurons, is necessary for a proper differentiation and is maintained in midbrain DA neurons in the adult brain (Perlmann and Wallén-Mackenzie 2004; Zetterström et al. 1997). Nurr1 null mice die within 24h after birth and conditional knockout of *Nr4a2* in DA neurons in adult animals caused decreased expression of DA markers, including *Th* and *Slc6a3* (DAT), followed by progressive loss of DA (Castillo et al. 1998; Kadkhodaei et al. 2009, 2013). Six additional DA-related transcription factors are downregulated in SNc (*Foxa2*, *Gata3*, *Pax5*, *Pitx3*, *En1* and *En2)*, but only *Pitx3* and *En1* are also downregulated in VTA of DAT:TH KO^Low^ animals. Together, our data suggests that DA is required for maintained expression of the DA-related genes required for DA biosynthesis and DA signaling by a novel regulatory loop with different susceptibility in SNc and VTA.

The majority of genes upregulated in DAT:TH KO VTA are annotated with GO terms connected to the regulation of trans-synaptic signaling and function of glutamatergic synapses. Reciprocal signaling between DA and glutamate is well documented. Decreased levels of midbrain DA resulting from unilateral 6-OHDA lesions, bilaterally increases levels of glutamate in striatum and rostromedial tegmental nucleus, with parallel changes in expression of glutamate receptors (Chang et al. 2019; Lindefors and Ungerstedt 1990; Rodríguez-Puertas et al. 1999). There are additional levels of crosstalk between glutamate and dopamine-glutamate neurons and their local synapses within the midbrain DA regions (Dobi et al. 2010; Morales and Root 2014; Viereckel et al. 2016; Wang and Morales 2009; Yamaguchi et al. 2015). In our study we detect increased levels of metabotropic glutamate receptor 1 and ionotropic glutamate receptor NMDA 3A. Increased glutamate signaling may play a role in a tuning of the VTA, since activation of glutamate receptors modulates firing of dopaminergic neurons and regulates extracellular DA levels (Ferrada et al. 2017; Floresco et al. 2003; Lodge and Grace 2006; Oakman et al. 1995).

Here we show data that suggests that DA is required for maintained expression of dopaminergic genes in SNc, and to a lesser degree in VTA, suggesting the existence of a feedback loop responsible for the maintenance of DA signaling. Furthermore, our study implicates an unexpected diversity in responses to DA loss between individual animals. Future studies should show whether the unusual class of animals expressing near-normal levels of the FLOXed Th mRNA represent a compensatory response, partly compensating for the DA loss.

## Acknowledgements

The laboratory of J.H. was supported by NIH 5R01GM084128. The authors state no conflict of interest.

## Author Contributions

All authors had full access to all the data in the study and take responsibility for the integrity of the data and the accuracy of the data analysis. Conceptualization, M.K., K.C., M.D., J.H.; Methodology, M.K., K.C., M.D., J.H.; Investigation, M.K., K.C., J.T.G, M.D.; Formal Analysis, M.K., K.C.; Resources: J.T.G, M.D., J.H.; Data curation, M.K.; Writing – Original Draft, M.K., J.H.; Writing – Review & Editing, M.K., K.C., M.D., J.H.; Visualization, M.K.; Supervision, M.D., J.H.; Funding Acquisition, M.D., J.H.

---

**Supplementary Figure 1.**
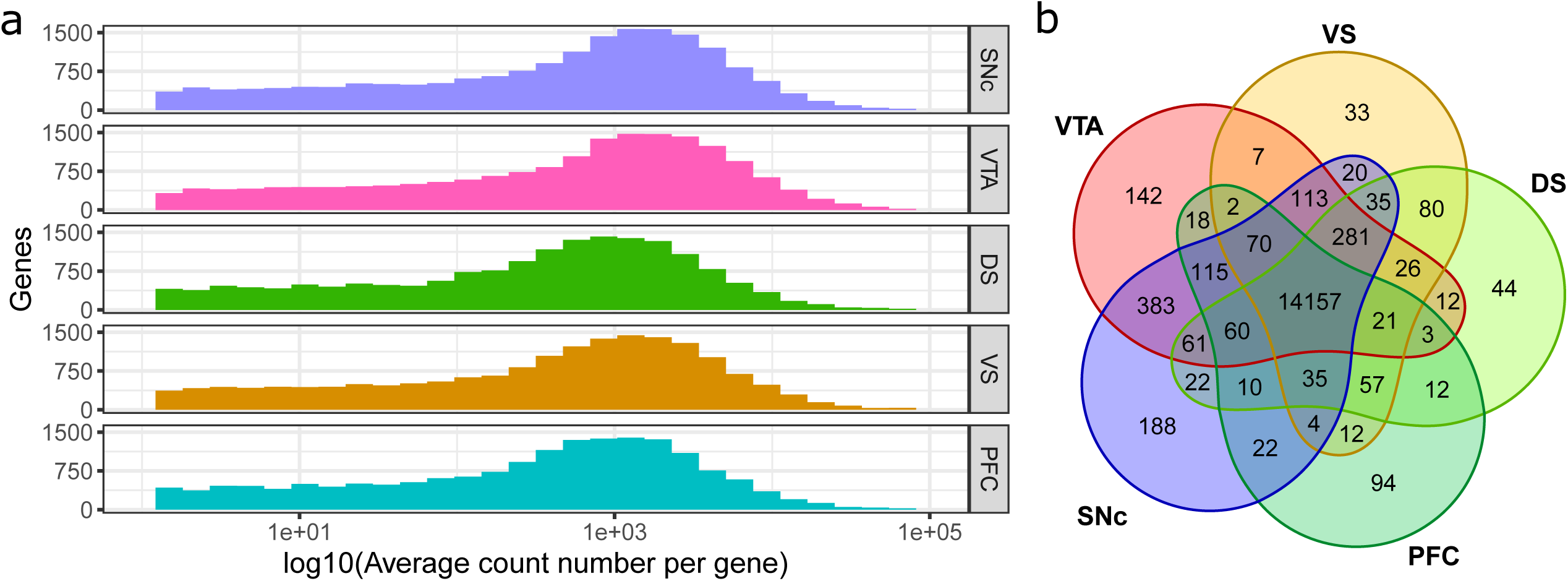
RNA-Seq Samples profiles. Input data for all analyses are DESeq2 normalized count values based on featureCounts output. A) Histograms representing the distributions of uniquely mapped reads per gene in examined brain regions. B) Venn diagram showing numbers of genes detected uniquely in the individual brain regions or in the intersections of two or more regions. Genes represented by fewer than 30 reads total were filtered out.

## Supplemental Information

**SUPPLEMENTAL FILE 1:** Spreadsheet with the detailed results of DE analysis.

**SUPPLEMENTAL FILE 2:** Spreadsheet with the results of GO analysis.

## References

Alessi, Amy et al. 2006. “γ-Syntrophin Scaffolding Is Spatially and Functionally Distinct from That of the α/β Syntrophins.” Experimental Cell Research 312(16): 3084–95.

Baertling, Fabian et al. 2017. “NDUFAF4 Variants Are Associated with Leigh Syndrome and Cause a Specific Mitochondrial Complex i Assembly Defect.” European Journal of Human Genetics 25(11): 1273–77.

Ballanger, Benedicte et al. 2016. “Imaging Dopamine and Serotonin Systems on MPTP Monkeys: A Longitudinal PET Investigation of Compensatory Mechanisms.” The Journal of Neuroscience 36(5): 1577–89. http://www.jneurosci.org/lookup/doi/10.1523/JNEUROSCI.2010-15.2016.

Baunez, C, and T W Robbins. 1999. “Effects of Dopamine Depletion of the Dorsal Striatum and Further Interaction with Subthalamic Nucleus Lesions in an Attentional Task in the Rat.” Neuroscience 92(4): 1343–56. http://www.ncbi.nlm.nih.gov/pubmed/10426489.

Bezard, E, and C E Gross. 1998. “Compensatory Mechanisms in Experimental and Human Parkinsonism: Towards a Dynamic Approach.” Progress in neurobiology 55(2): 93–116. http://www.ncbi.nlm.nih.gov/pubmed/9618745.

Bezard, Erwan et al. 2001. “Upregulation of Striatal Preproenkephalin Gene Expression Occurs before the Appearance of Parkinsonian Signs in 1-Methyl-4-Phenyl- 1,2,3,6- Tetrahydropyridine Monkeys.” Neurobiology of Disease 8(2): 343–50. https://linkinghub.elsevier.com/retrieve/pii/S0969996100903759.

Bezard, Erwan, Christian E. Gross, and Jonathan M. Brotchie. 2003. “Presymptomatic Compensation in Parkinson’s Disease Is Not Dopamine-Mediated.” Trends in Neurosciences 26(4): 215–21. https://linkinghub.elsevier.com/retrieve/pii/S0166223603000389.

Björklund, Anders, and Stephen B. Dunnett. 2007. “Dopamine Neuron Systems in the Brain: An Update.” Trends in Neurosciences 30(5): 194–202. https://linkinghub.elsevier.com/retrieve/pii/S0166223607000677.

Blesa, J. et al. 2012. “The Nigrostriatal System in the Presymptomatic and Symptomatic Stages in the MPTP Monkey Model: A PET, Histological and Biochemical Study.” Neurobiology of Disease 48(1): 79–91. https://linkinghub.elsevier.com/retrieve/pii/S0969996112002045.

Bourens, Myriam, Aren Boulet, Scot C. Leary, and Antoni Barrientos. 2014. “Human COX20 Cooperates with SCO1 and SCO2 to Mature COX2 and Promote the Assembly of Cytochrome c Oxidase.” Human Molecular Genetics 23(11): 2901–13.

Brichta, Lars, and Paul Greengard. 2014. “Molecular Determinants of Selective Dopaminergic Vulnerability in Parkinsonâ€TMs Disease: An Update.” Frontiers in Neuroanatomy 8(December): 1–16. http://journal.frontiersin.org/article/10.3389/fnana.2014.00152/abstract.

Carey, A. N. et al. 2009. “Endogenous Opioid Activation Mediates Stress-Induced Deficits in Learning and Memory.” Journal of Neuroscience 29(13): 4293–4300.

Castillo, Susan O. et al. 1998. “Dopamine Biosynthesis Is Selectively Abolished in Substantia Nigra/Ventral Tegmental Area but Not in Hypothalamic Neurons in Mice with Targeted Disruption of the Nurr1 Gene.” Molecular and Cellular Neuroscience 11(1–2): 36–46. https://linkinghub.elsevier.com/retrieve/pii/S104474319890673X.

Chang, Yongli et al. 2019. “Enhanced AMPA Receptor-Mediated Excitatory Transmission in the Rodent Rostromedial Tegmental Nucleus Following Lesion of the Nigrostriatal Pathway.” Neurochemistry International 122(September 2018): 85–93. https://doi.org/10.1016/j.neuint.2018.11.007.

Chen, L. et al. 2008. “Unregulated Cytosolic Dopamine Causes Neurodegeneration Associated with Oxidative Stress in Mice.” Journal of Neuroscience 28(2): 425–33.

Chin, Mark H. et al. 2008. “Mitochondrial Dysfunction, Oxidative Stress, and Apoptosis Revealed by Proteomic and Transcriptomic Analyses of the Striata in Two Mouse Models of Parkinson’s Disease.” Journal of Proteome Research 7(2): 666–77. http://pubs.acs.org/doi/abs/10.1021/pr070546l.

Chudasama, Y, and T W Robbins. 2006. “Functions of Frontostriatal Systems in Cognition: Comparative Neuropsychopharmacological Studies in Rats, Monkeys and Humans.” Biological psychology 73(1): 19–38. http://www.ncbi.nlm.nih.gov/pubmed/16546312.

Cools, R, R A Barker, B J Sahakian, and T W Robbins. 2001. “Enhanced or Impaired Cognitive Function in Parkinson’s Disease as a Function of Dopaminergic Medication and Task Demands.” Cerebral cortex (New York, N.Y. : 1991) 11(12): 1136–43. http://www.ncbi.nlm.nih.gov/pubmed/11709484.

Cools, Roshan, Roger A. Barker, Barbara J. Sahakian, and Trevor W. Robbins. 2003. “L-Dopa Medication Remediates Cognitive Inflexibility, but Increases Impulsivity in Patients with Parkinson’s Disease.” Neuropsychologia 41(11): 1431–41.

Da Cunha, Claudio et al. 2002. “The Lesion of the Rat Substantia Nigra Pars Compacta Dopaminergic Neurons as a Model for Parkinson’s Disease Memory Disabilities.” Cellular and molecular neurobiology 22(3): 227–37. http://www.ncbi.nlm.nih.gov/pubmed/12469866.

Darvas, Martin, Charles W. Henschen, and Richard D. Palmiter. 2014. “Contributions of Signaling by Dopamine Neurons in Dorsal Striatum to Cognitive Behaviors Corresponding to Those Observed in Parkinson’s Disease.” Neurobiology of disease 65(11): 112–23. http://www.ncbi.nlm.nih.gov/pubmed/23959937.

Darvas, Martin, and Richard D Palmiter. 2009. “Restriction of Dopamine Signaling to the Dorsolateral Striatum Is Sufficient for Many Cognitive Behaviors.” Proceedings of the National Academy of Sciences of the United States of America 106(34): 14664–69. http://www.ncbi.nlm.nih.gov/pubmed/19667174.

Darvas, Martin, and Richard D Palmiter. 2011. “Contributions of Striatal Dopamine Signaling to the Modulation of Cognitive Flexibility.” Biological psychiatry 69(7): 704–7. http://www.ncbi.nlm.nih.gov/pubmed/21074144.

Darvas, Martin, Amanda M Wunsch, Jeffrey T Gibbs, and Richard D Palmiter. 2014. “Dopamine Dependency for Acquisition and Performance of Pavlovian Conditioned Response.” Proceedings of the National Academy of Sciences of the United States of America 111(7): 2764–69. http://www.ncbi.nlm.nih.gov/pubmed/24550305.

Dawson, Ted M., and Valina L. Dawson. 2003. “Molecular Pathways of Neurodegeneration in Parkinson’s Disease.” Science 302(5646): 819–22.

Dobi, A. et al. 2010. “Glutamatergic and Nonglutamatergic Neurons of the Ventral Tegmental Area Establish Local Synaptic Contacts with Dopaminergic and Nondopaminergic Neurons.” Journal of Neuroscience 30(1): 218–29. http://doi.wiley.com/10.1111/j.1460-9568.2008.06576.x.

Domanskyi, Andrii et al. 2014. “Transcription Factors Foxa1 and Foxa2 Are Required for Adult Dopamine Neurons Maintenance.” Frontiers in Cellular Neuroscience 8(September): 1–11. http://journal.frontiersin.org/article/10.3389/fncel.2014.00275/abstract.

Elliott, Leah E., Scott A. Saracco, and Thomas D. Fox. 2012. “Multiple Roles of the Cox20 Chaperone in Assembly of Saccharomyces Cerevisiae Cytochrome C Oxidase.” Genetics 190(2): 559–67.

Elmore, Susan. 2007. “Apoptosis: A Review of Programmed Cell Death.” Toxicologic Pathology 35(4): 495–516. http://www.ncbi.nlm.nih.gov/pubmed/21959306.

Ferrada, Carla, Ramón Sotomayor-Zárate, Jorge Abarca, and Katia Gysling. 2017. “The Activation of Metabotropic Glutamate 5 Receptors in the Rat Ventral Tegmental Area Increases Dopamine Extracellular Levels.” NeuroReport 28(1): 28–34.

Floresco, Stan B. et al. 2003. “Afferent Modulation of Dopamine Neuron Firing Differentially Regulates Tonic and Phasic Dopamine Transmission.” Nature Neuroscience 6(9): 968–73.

Gao, X. et al. 2008. “Gene-Gene Interaction between FGF20 and MAOB in Parkinson Disease.” Annals of Human Genetics 72(2): 157–62.

Garrett Morgan, R. et al. 2015. “Relative Contributions of Severe Dopaminergic Neuron Ablation and Dopamine Depletion to Cognitive Impairment.” Experimental Neurology 271(11): 205–14. https://linkinghub.elsevier.com/retrieve/pii/S0014488615300236.

Gerfen, Charles R, and Heinz Steiner. 1998. “Role of Dynorphin and Enkephalin in the Regulation of Striatal Output Pathways and Behavior.” Experimental Brain Research 123(1): 60–76. http://www.springerlink.com/index/CKAJM5L3X9WYM1PN.pdf.

Golden, Judith P. et al. 2013. “Dopamine-Dependent Compensation Maintains Motor Behavior in Mice with Developmental Ablation of Dopaminergic Neurons.” The Journal of Neuroscience 33(43): 17095–107. http://www.jneurosci.org/lookup/doi/10.1523/JNEUROSCI.0890-13.2013.

Grimes, David A. et al. 2006. “Translated Mutation in the Nurr1 Gene as a Cause for Parkinson’s Disease.” Movement Disorders 21(7): 906–9.

Gubellini, P., and P. Kachidian. 2015. “Animal Models of Parkinson’s Disease: An Updated Overview.” Revue Neurologique 171(11): 750–61. http://dx.doi.org/10.1016/j.neurol.2015.07.011.

Gubellini, Paolo, Barbara Picconi, Massimiliano Di Filippo, and Paolo Calabresi. 2010. “Downstream Mechanisms Triggered by Mitochondrial Dysfunction in the Basal Ganglia: From Experimental Models to Neurodegenerative Diseases.” Biochimica et Biophysica Acta - Molecular Basis of Disease 1802(1): 151–61. http://dx.doi.org/10.1016/j.bbadis.2009.08.001.

Halliday, Glenda, Mariese Hely, Wayne Reid, and John Morris. 2008. “The Progression of Pathology in Longitudinally Followed Patients with Parkinson’s Disease.” Acta Neuropathologica 115(4): 409–15.

Haugas, Maarja et al. 2016. “Gata2 and Gata3 Regulate the Differentiation of Serotonergic and Glutamatergic Neuron Subtypes of the Dorsal Raphe.” Development 143(23): 4495–4508. http://dev.biologists.org/lookup/doi/10.1242/dev.136614.

Hayashi, Takaharu et al. 2015. “Higd1a Is a Positive Regulator of Cytochrome c Oxidase.” Proceedings of the National Academy of Sciences 112(5): 1553–58.

Henschen, Charles W., Richard D. Palmiter, and Martin Darvas. 2013. “Restoration of Dopamine Signaling to the Dorsal Striatum Is Sufficient for Aspects of Active Maternal Behavior in Female Mice.” Endocrinology 154(11): 4316–27. http://www.ncbi.nlm.nih.gov/pubmed/23959937.

Hornykiewicz, O. 1975. “Brain Monoamines and Parkinsonism.” National Institute on Drug Abuse research monograph series (3): 13–21. http://www.ncbi.nlm.nih.gov/pubmed/787796.

Jackson-Lewis, Vernice, Michael Jakowec, Robert E Burke, and Serge Przedborski. 1995. “Time Course and Morphology of Dopaminergic Neuronal Death Caused by the Neurotoxin 1-Methyl-4-Phenyl-1,2,3,6-Tetrahydropyridine.” Neurodegeneration 4(3): 257–69. c:I.

Jackson, Chad R et al. 2012. “Retinal Dopamine Mediates Multiple Dimensions of Light-Adapted Vision.” The Journal of neuroscience : the official journal of the Society for Neuroscience 32(27): 9359–68. http://www.ncbi.nlm.nih.gov/pubmed/22764243.

Jacobs, F. M. J. et al. 2011. “Retinoic Acid-Dependent and -Independent Gene-Regulatory Pathways of Pitx3 in Meso-Diencephalic Dopaminergic Neurons.” Development 138(23): 5213–22. http://dev.biologists.org/cgi/doi/10.1242/dev.071704.

Jiang, Min-Yao et al. 2017. “Mitochondrion-Associated Protein Peroxiredoxin 3 Promotes Benign Prostatic Hyperplasia through Autophagy Suppression and Pyroptosis Activation.” Oncotarget 8(46): 80295–302.

Jorge-Finnigan, Ana et al. 2017. “Phosphorylation at Serine 31 Targets Tyrosine Hydroxylase to Vesicles for Transport along Microtubules.” Journal of Biological Chemistry 292(34): 14092–107.

Kadkhodaei, B. et al. 2009. “Nurr1 Is Required for Maintenance of Maturing and Adult Midbrain Dopamine Neurons.” Journal of Neuroscience 29(50): 15923–32. http://www.jneurosci.org/cgi/doi/10.1523/JNEUROSCI.3910-09.2009.

Kadkhodaei, B. 2013. “Transcription Factor Nurr1 Maintains Fiber Integrity and Nuclear-Encoded Mitochondrial Gene Expression in Dopamine Neurons.” Proceedings of the National Academy of Sciences 110(6): 2360–65. http://www.pnas.org/cgi/doi/10.1073/pnas.1221077110.

Kalia, Lorraine V., and Anthony E. Lang. 2015. “Parkinson’s Disease.” The Lancet 386(9996): 896–912. http://dx.doi.org/10.1016/S0140-6736(14)61393-3.

Kato, Hiroki, and Katsuyoshi Mihara. 2008. “Identification of Tom5 and Tom6 in the Preprotein Translocase Complex of Human Mitochondrial Outer Membrane.” Biochemical and Biophysical Research Communications 369(3): 958–63.

Kawazoe, Tomoya et al. 2007. “Structural Basis of D-DOPA Oxidation by d-Amino Acid Oxidase: Alternative Pathway for Dopamine Biosynthesis.” Biochemical and Biophysical Research Communications 355(2): 385–91.

Kim, Daehwan, Ben Langmead, and Steven L. Salzberg. 2015. “HISAT: A Fast Spliced Aligner with Low Memory Requirements.” Nature Methods 12(4): 357–60.

Kim, J.-I. et al. 2015. “Aldehyde Dehydrogenase 1a1 Mediates a GABA Synthesis Pathway in Midbrain Dopaminergic Neurons.” Science 350(6256): 102–6. http://www.sciencemag.org/cgi/doi/10.1126/science.aac4690.

Koranda, Jessica L et al. 2014. “Nicotinic Receptors Regulate the Dynamic Range of Dopamine Release in Vivo.” Journal of Neurophysiology 111: 103–11. http://www.ncbi.nlm.nih.gov/pubmed/24089398.

Larhammar, Martin et al. 2015. “SLC10A4 Is a Vesicular Amine-Associated Transporter Modulating Dopamine Homeostasis.” Biological Psychiatry 77(6): 526–36. http://dx.doi.org/10.1016/j.biopsych.2014.07.017.

Lautenschläger, Janin et al. 2018. “An Easy-to-Implement Protocol for Preparing Postnatal Ventral Mesencephalic Cultures.” Frontiers in Cellular Neuroscience 12(March): 1–10.

Le, Wei dong et al. 2003. “Mutations in NR4A2 Associated with Familial Parkinson Disease.” Nature Genetics 33(1): 85–89.

Leino, Sakari et al. 2018. “Attenuated Dopaminergic Neurodegeneration and Motor Dysfunction in Hemiparkinsonian Mice Lacking the Α Nicotinic Acetylcholine Receptor Subunit.” Neuropharmacology 138: 371–80.

De Leonibus, Elvira et al. 2007. “Spatial Deficits in a Mouse Model of Parkinson Disease.” Psychopharmacology 194(4): 517–25. http://www.ncbi.nlm.nih.gov/pubmed/17619858.

Liao, Yang, Gordon K. Smyth, and Wei Shi. 2014. “FeatureCounts: An Efficient General Purpose Program for Assigning Sequence Reads to Genomic Features.” Bioinformatics 30(7): 923–30.

Lindefors, Nils, and Urban Ungerstedt. 1990. “Bilateral Regulation of Glutamate Tissue and Extracellular Levels in Caudate-Putamen by Midbrain Dopamine Neurons.” Neuroscience Letters 115(2–3): 248–52.

Liu, Zhilei et al. 2016. “Hydrogen Peroxide Mediated Mitochondrial UNG1-PRDX3 Interaction and UNG1 Degradation.” Free Radical Biology and Medicine 99: 54–62. http://dx.doi.org/10.1016/j.freeradbiomed.2016.07.030.

Lodge, D. J., and A. A. Grace. 2006. “The Laterodorsal Tegmentum Is Essential for Burst Firing of Ventral Tegmental Area Dopamine Neurons.” Proceedings of the National Academy of Sciences 103(13): 5167–72.

Love, Michael I, Wolfgang Huber, and Simon Anders. 2014. “Moderated Estimation of Fold Change and Dispersion for RNA-Seq Data with DESeq2.” : 1–21.

Luk, Kelvin C et al. 2012. “Pathological α-Synuclein Transmission Initiates Parkinson-like Neurodegeneration in Nontransgenic Mice.” Science (New York, N.Y.) 338(6109): 949–53. http://www.ncbi.nlm.nih.gov/pubmed/23161999.

Lundblad, Martin, Mickael Decressac, Bengt Mattsson, and Anders Björklund. 2012. “Impaired Neurotransmission Caused by Overexpression of α-Synuclein in Nigral Dopamine Neurons.” Proceedings of the National Academy of Sciences of the United States of America 109(9): 3213–19. http://www.ncbi.nlm.nih.gov/pubmed/22315428.

Marcus, Dana, Michal Lichtenstein, Ann Saada, and Haya Lorberboum-Galski. 2013. “Replacement of the C6ORF66 Assembly Factor (NDUFAF4) Restores Complex I Activity in Patient Cells.” Molecular Medicine 19(1): 124–34.

Melief, E.J. et al. 2016. “Characterization of Cognitive Impairments and Neurotransmitter Changes in a Novel Transgenic Mouse Lacking Slc10a4.” Neuroscience 324(5): 399–406. https://linkinghub.elsevier.com/retrieve/pii/S0306452216300070.

Melief, Erica J. et al. 2018. “Loss of Glutamate Signaling from the Thalamus to Dorsal Striatum Impairs Motor Function and Slows the Execution of Learned Behaviors.” npj Parkinson’s Disease 4(1). http://dx.doi.org/10.1038/s41531-018-0060-6.

Morales, M., and D. H. Root. 2014. “Glutamate Neurons within the Midbrain Dopamine Regions.” Neuroscience 282: 60–68. http://dx.doi.org/10.1016/j.neuroscience.2014.05.032.

Moriguchi, T. 2006. “Gata3 Participates in a Complex Transcriptional Feedback Network to Regulate Sympathoadrenal Differentiation.” Development 133(19): 3871–81. http://dev.biologists.org/cgi/doi/10.1242/dev.02553.

Mura, Anna, and Joram Feldon. 2003. “Spatial Learning in Rats Is Impaired after Degeneration of the Nigrostriatal Dopaminergic System.” Movement disorders : official journal of the Movement Disorder Society 18(8): 860–71. http://www.ncbi.nlm.nih.gov/pubmed/12889075.

Nakashima, Akira et al. 2005. “Deletion of N-Terminus of Human Tyrosine Hydroxylase Type 1 Enhances Stability of the Enzyme in AtT-20 Cells.” 120: 110–20.

Oakman, SA et al. 1995. “Distribution of Pontomesencephalic Cholinergic Neurons Projecting to Substantia Nigra Differs Significantly from Those Projecting to Ventral Tegmental Area.” The Journal of Neuroscience 15(9): 5859–69. http://www.jneurosci.org/lookup/doi/10.1523/JNEUROSCI.15-09-05859.1995.

Ott, Christine et al. 2015. “Detailed Analysis of the Human Mitochondrial Contact Site Complex Indicate a Hierarchy of Subunits.” PLoS ONE 10(3): 1–15.

Patel, Suman, Kavita Singh, Seema Singh, and Mahendra Pratap Singh. 2008. “Gene Expression Profiles of Mouse Striatum in Control and Maneb + Paraquat-Induced Parkinson’s Disease Phenotype: Validation of Differentially Expressed Energy Metabolizing Transcripts.” Molecular Biotechnology 40(1): 59–68.

Pattarini, R., Y. Rong, C. Qu, and J. I. Morgan. 2008. “Distinct Mechanisms of 1-Methyl-4-Phenyl-1,2,3,6-Tetrahydropyrimidine Resistance Revealed by Transcriptome Mapping in Mouse Striatum.” Neuroscience 155(4): 1174–94.

Perez, Xiomara A. et al. 2008. “Pre-Synaptic Dopaminergic Compensation after Moderate Nigrostriatal Damage in Non-Human Primates.” Journal of Neurochemistry 105(5): 1861–72. http://doi.wiley.com/10.1111/j.1471-4159.2008.05268.x.

Perlmann, Thomas, and Asa Wallén-Mackenzie. 2004. “Nurr1, an Orphan Nuclear Receptor with Essential Functions in Developing Dopamine Cells.” Cell and tissue research 318(1): 45–52. http://link.springer.com/10.1007/s00441-004-0974-7.

Piluso, Giulio et al. 2000. “Γ1- and Γ2-Syntrophins, Two Novel Dystrophin-Binding Proteins Localized in Neuronal Cells.” Journal of Biological Chemistry 275(21): 15851–60.

Postuma, Ronald B. et al. 2012. “Identifying Prodromal Parkinson’s Disease: Pre-Motor Disorders in Parkinson’s Disease.” Movement Disorders 27(5): 617–26.

Quik, Maryka, and Susan Wonnacott. 2011. “6 2* and 4 2* Nicotinic Acetylcholine Receptors As Drug Targets for Parkinson’s Disease.” Pharmacological Reviews 63(4): 938–66. http://pharmrev.aspetjournals.org/cgi/doi/10.1124/pr.110.003269.

Reid, M S et al. 1990. “Striatonigral GABA, Dynorphin, Substance P and Neurokinin A Modulation of Nigrostriatal Dopamine Release: Evidence for Direct Regulatory Mechanisms.” Experimental brain research 82(2): 293–303. http://www.ncbi.nlm.nih.gov/pubmed/1704847.

Rodríguez-Puertas, R., M. Herrera-Marschitz, J. Koistinaho, and T. Hökfelt. 1999. “Dopamine D1 Receptor Modulation of Glutamate Receptor Messenger RNA Levels in the Neocortex and Neostriatum of Unilaterally 6-Hydroxydopamine-Lesioned Rats.” Neuroscience 89(3): 781–97.

S., Andrews. 2010. “FastQC: A Quality Control Tool for High Throughput Sequence Data.” http://www.bioinformatics.babraham.ac.uk/projects/fastqc.

Saada, Ann et al. 2009. “Mutations in NDUFAF3 (C3ORF60), Encoding an NDUFAF4 (C6ORF66)-Interacting Complex I Assembly Protein, Cause Fatal Neonatal Mitochondrial Disease.” American Journal of Human Genetics 84(6): 718–27.

Sawamoto, Nobukatsu et al. 2008. “Cognitive Deficits and Striato-Frontal Dopamine Release in Parkinson’s Disease.” Brain 131(5): 1294–1302.

Simon, Horst H et al. 2003. “Midbrain Dopaminergic Neurons: Determination of Their Developmental Fate by Transcription Factors.” Annals of the New York Academy of Sciences 991: 36–47.

Simunovic, Filip et al. 2009. “Gene Expression Profiling of Substantia Nigra Dopamine Neurons: Further Insights into Parkinson’s Disease Pathology.” Brain 132(7): 1795–1809.

Singh, Kavita et al. 2010. “Nicotine- and Caffeine-Mediated Changes in Gene Expression Patterns of MPTP-Lesioned Mouse Striatum: Implications in Neuroprotection Mechanism.” Chemico-Biological Interactions 185(2): 81–93. http://dx.doi.org/10.1016/j.cbi.2010.03.015.

Sivam, S. P. 1996. “Dopaminergic Regulation of Postnatal Development of Dynorphin Neurons in Rat Striatum.” Neuropeptides 30(1): 103–7.

Sleeman, Isobel J., Eugene L. Boshoff, and Susan Duty. 2012. “Fibroblast Growth Factor-20 Protects against Dopamine Neuron Loss in Vitro and Provides Functional Protection in the 6-Hydroxydopamine-Lesioned Rat Model of Parkinson’s Disease.” Neuropharmacology 63(7): 1268–77. http://dx.doi.org/10.1016/j.neuropharm.2012.07.029.

Smith, M. P., and W. A. Cass. 2007. “Oxidative Stress and Dopamine Depletion in an Intrastriatal 6-Hydroxydopamine Model of Parkinson’s Disease.” Neuroscience 144(3): 1057–66.

Sulzer, David, Stephanie J. Cragg, and Margaret E. Rice. 2016. “Striatal Dopamine Neurotransmission: Regulation of Release and Uptake.” Basal Ganglia 6(3): 123–48. http://dx.doi.org/10.1016/j.baga.2016.02.001.

Tekin, Izel, Robert Roskoski, Nurgul Carkaci-Salli, and Kent E. Vrana. 2014. 121 Journal of Neural Transmission Complex Molecular Regulation of Tyrosine Hydroxylase.

Triepels, R. et al. 1999. “The Human Nuclear-Encoded Acyl Carrier Subunit (NDUFAB1) of the Mitochondrial Complex I in Human Pathology.” Journal of Inherited Metabolic Disease 22(2): 163–73.

Ungerstedt, Urban. 1968. “6-Ohda Induced Degeneration of Central Monoamine Neurons (Ungerstedt 1968).Pdf.” 5: 107–10.

Ungerstedt, Urban, and Gordon W Arbuthnott. 1970. “Quantitative Recording of Rotational Behavior in Rats after 6-Hydroxy-Dopamine Lesions of the Nigrostriatal Dopamine System.” Brain Research 24(3): 485–93. http://linkinghub.elsevier.com/retrieve/pii/0006899370901873.

Viereckel, Thomas et al. 2016. “Midbrain Gene Screening Identifies a New Mesoaccumbal Glutamatergic Pathway and a Marker for Dopamine Cells Neuroprotected in Parkinson’s Disease.” Scientific Reports 6(September): 1–16. http://dx.doi.org/10.1038/srep35203.

Van Der Walt, Joelle M et al. 2004. “Fibroblast Growth Factor 20 Polymorphisms and Haplotypes Strongly Influence Risk of Parkinson Disease.” Am. J. Hum. Genet 74: 1121–27. https://ac.els-cdn.com/S0002929707628390/1-s2.0-S0002929707628390-main.pdf?_tid=587fff88-e776-4b11-8152-0f2b1638702f&acdnat=1537861235_b3f521df7fa23a4d471224ace29797d7.

Wang, Hui-Ling, and Marisela Morales. 2009. “Pedunculopontine and Laterodorsal Tegmental Nuclei Contain Distinct Populations of Cholinergic, Glutamatergic and GABAergic Neurons in the Rat.” European Journal of Neuroscience 29(2): 340–58. http://doi.wiley.com/10.1111/j.1460-9568.2008.06576.x.

Wise, Roy A. 2004. “Dopamine, Learning and Motivation.” Nature Reviews Neuroscience 5(6): 483–94. http://www.nature.com/articles/nrn1406.

Xu, Ming et al. 1994. “Dopamine D1 Receptor Mutant Mice Are Deficient in Striatal Expression of Dynorphin and in Dopamine-Mediated Behavioral Responses.” Cell 79(4): 729–42.

Xu, P. Y. et al. 2002. “Association of Homozygous 7048G7049 Variant in the Intron Six of Nurr1 Gene with Parkinson’s Disease.” Neurology 58(6): 881–84.

Yamaguchi, Tsuyoshi et al. 2015. “Glutamatergic and Dopaminergic Neurons in the Mouse Ventral Tegmental Area.” European Journal of Neuroscience 41(6): 760–72.

You, Zhi-bing, Mario Herrera-marschitz, and Lars Terenius. 1999. “Modulation of Neurotransmitter Release in the Basal Ganglia of the Rat Brain by Dynorphin Peptides 1.” Journal of Pharmacology and Experimental Therapeutics 290(3): 1307–15. http://jpet.aspetjournals.org/content/290/3/1307.abstract.

Zetterström, R H et al. 1997. “Dopamine Neuron Agenesis in Nurr1-Deficient Mice.” Science (New York, N.Y.) 276(5310): 248–50. http://www.ncbi.nlm.nih.gov/pubmed/9092472.

Zheng, Kangni, Bobak Heydari, and David K Simon. 2003. “A Common NURR1 Polymorphism Associated with Parkinson Disease and Diffuse Lewy Body Disease.” Archives of neurology 60(5): 722–25. http://www.ncbi.nlm.nih.gov/pubmed/12756136.

Zhou, Qun Yong, and Richard D. Palmiter. 1995. “Dopamine-Deficient Mice Are Severely Hypoactive, Adipsic, and Aphagic.” Cell 83(7): 1197–1209.

Zhuang, Xiaoxi et al. 2005. “Targeted Gene Expression in Dopamine and Serotonin Neurons of the Mouse Brain.” Journal of neuroscience methods 143(1): 27–32. http://www.ncbi.nlm.nih.gov/pubmed/15763133.

Zurita Rendón, Olga, and Eric A. Shoubridge. 2012. “Early Complex I Assembly Defects Result in Rapid Turnover of the ND1 Subunit.” Human Molecular Genetics 21(17): 3815–24.

